# Discovery of Novel, Potent and Orally Bioavailable SMARCA2 PROTACs with Synergistic Anti-tumor Activity in Combination with KRAS G12C Inhibitors

**DOI:** 10.1101/2024.08.23.608456

**Authors:** Sasikumar Kotagiri, Yawen Wang, Yanyan Han, Xiaobing Liang, Nicholas Blazanin, Phuong Kieu Nguyen, Yongying Jiang, Yonathan Lissanu

## Abstract

Cancer genomic studies have identified frequent mutations in subunits of the SWI/SNF chromatin remodeling complex including *SMARCA4* in non-small cell lung cancer with a frequency of up to 33% in advanced stage disease, making it the most frequently mutated complex in lung cancer. We and others have identified *SMARCA2* to be synthetic lethal to *SMARCA4,* indicating SMARCA2 is a high value therapeutic target. Here, we disclose the discovery and characterization of potent, selective and orally bioavailable Cereblon-based SMARCA2 PROTACs. Biochemically, YDR1 and YD54 are potent SMARCA2 degraders with an average DC_50_ of 7.7nM and 3.5nM respectively in *SMARCA4* mutant lung cancer cells. Phenotypically, both YDR1 and YD54 selectively inhibited growth of *SMARCA4* mutant cancer cells. Further, we showed anti-tumor growth inhibitory activity of YDR1 and YD54 in *SMARCA4* mutant xenograft models of lung cancer. Finally, we show that YDR1 and YD54 synergize with the KRAS G12C inhibitor sotorasib to inhibit growth of *SMARCA4* and *KRAS G12C* co-mutant lung cancer cells. These findings provide additional evidence for the utility of single agent or combination regimens containing SMARCA2 PROTACs as synthetic lethal therapeutics against *SMARCA4* mutant cancers.

## INTRODUCTION

A major discovery of large-scale cancer genomic studies is the identification of frequent mutations in subunits of the SWI/SNF (SWItch/Sucrose Non-Fermenting) chromatin remodeling complex including *SMARCA4* (also known as *BRG1*) and *ARID1A* ranging from 16% in early stage disease to 33% in advanced lung cancer^1–7^. Additionally, a meta-analysis of 44 cancer genomic studies has shown that 20% of all solid tumors have mutations in subunits of the SWI/SNF complex making it one of the most frequently mutated complexes in cancer^6^. The SWI/SNF complex is a large multi-subunit complex that uses ATP hydrolysis to remodel nucleosomes and enable chromatin dependent processes such as transcription, DNA repair and replication ^8, 9^. SMARCA2 (also known as BRM) and SMARCA4 are mutually exclusive core catalytic subunits with ATPase enzymatic activity. Mutations in *SMARCA4* are inactivating and cannot be directly targeted therapeutically. Thus, there is intense effort to identify synthetic lethal or other vulnerabilities in *SMARCA4* mutants. While a number of studies have reported enhanced sensitivity of *SMARCA4*-mutant lung cancers to inhibition of oxidative phosphorylation (OXPHOS)^10^, Aurora kinase A^11^, EZH2^12^, ATR^13^, and KDM6 methytransferases^14^, none of these have progressed into clinical success. Importantly, we and others have identified that the paralog *SMARCA2* shows a synthetic lethal genetic interaction with *SMARCA4* in lung cancer cell lines suggesting it is a high value therapeutic target ^15–20^. SMARCA2 is composed of helicase-SANT-associated (HSA) domain, DEXDc and Helicase domains (that together constitute the ATPase domain), Snf2 ATP coupling (SnAC) domain and bromodomain. From these, the ATPase and bromodomains are readily amenable for small molecule inhibition. As anti-cancer agents, inhibitors targeting the bromodomain were found to be not efficacious, while those targeting the ATPase domain of SMARCA2 were not selective and associated with undesired off-target activities against other ATPases such as SMARCA4, highlighting the need for alternative approaches of targeting SMARCA2^16, 21^. Proteolysis targeting chimeras (PROTACs) have emerged as an innovative class of therapeutics to expand the repertoire of the targetable genome ^22^. PROTACs are heterobifunctional molecules that tether a small molecule ligand of a protein of interest to an E3 ubiquitin ligase binder and result in induced proximity of a target protein to the cellular ubiquitination machinery leading to ubiquitination and proteasomal degradation of target protein ^22^. Recently, several SMARCA2 PROTACs including ACBI2, A947 and SMD-3040 have been reported with varying pharmacokinetic properties and wide range of selectivity over SMARCA4^23–25^. While these developments are a major leap for the field, all these PROTACs utilize ligands to recruit the VHL E3 ubiquitin ligase. The only other PROTAC that targets SMARCA2 using Cereblon E3 ligase is a dual SMARCA4 and SMARCA2 degrader which actually has higher selectivity to SMARCA4^26^. So far, VHL and Cereblon are the most commonly utilized E3 ligases in PROTAC design. Importantly, Cereblon-based PROTACs have advanced the most in clinical development including ARV-471 in Phase 3 clinical trials for breast cancer^27^. Furthermore, acquired resistance to PROTACs in cancer treatment in model systems was primarily caused by genomic alterations that disable core components of the E3 ligase utilized^28^. Thus, having an array of PROTACs that use different E3 ligases for any therapeutic target is highly desirable. As all the above described selective SMARCA2 PROTACs were based on VHL ligase, we decided to pursue development of Cereblon-based SMARCA2 PROTACs.

Here, we report the discovery of YDR1 and YD54, potent, selective and orally bioavailable SMARCA2 degrading PROTACs based on the E3 ligase Cereblon by conjugating a SMARCA2 bromodomain ligand to pomalidomide (E3 ligase ligand) through rigid chemical linkers (Figure 1). We show that YDR1 and YD54 potently degrade SMARCA2 and selectively inhibit the growth of *SMARCA4*-mutant lung cancer cell lines. We characterized the microsomal stability and possible pathways of metabolic inactivation and importantly, we showed that YDR1 and YD54 are orally bioavailable with favorable pharmacokinetic properties for *in vivo* studies. We further showed that YDR1 and YD54 are potently active in mice and degrade SMARCA2 in various tissues and xenograft tumors. We also demonstrated that YDR1 and YD54 were well-tolerated in mice and able to inhibit growth of *SMARCA4* mutant xenograft tumors. Finally, we showed that YDR1 and YD54 have synergistic growth inhibitory effect in combination with the KRAS G12C inhibitor sotorasib in inhibiting growth of *SMARCA4* and *KRAS ^G12C^*co-mutant cells *in vitro* and xenograft tumors *in vivo*. To our knowledge, these are the first orally bioavailable Cereblon-based selective SMARCA2 PROTACs. In summary, this report provides novel SMARCA2 PROTACs as chemical probes to interrogate the *in vitro* and *in vivo* functions of SMARCA2 and serve as potential drug leads for *SMARCA4* mutant cancers.

**Figure 1:**
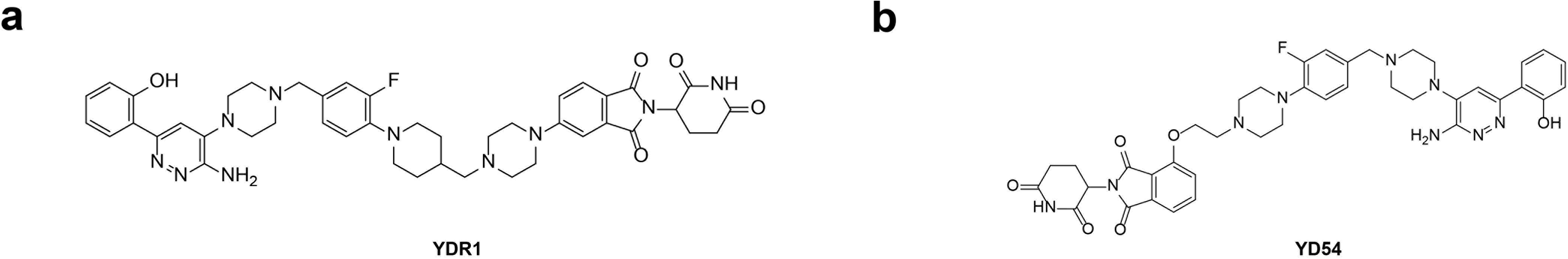
Structures of YDR1 and YD54. Chemical structures of YDR1 and YD54.

## RESULTS

### PROTAC Design

Heterobifunctional PROTACs targeted at SMARCA2 were designed based on the bromodomain binding ligand Gen-1 that was identified from a series of 2-(6-aminopyridazin-3-yl) phenols^17, 29^. Further, the compounds were based on pomalidomide and designed to recruit the E3 ubiquitin ligase cereblon (CRBN). We have synthesized heterobifunctional compounds we termed YDR1 and YD54 using variable fluorophenyl containing piperidinyl-piperazine or piperazine linkers respectively based on encouraging recent reports that such rigid linkers can help improve aqueous solubility, cell permeability and improve pharmacokinetic properties of the degrader (Figure 1)^30,31^.

### Chemistry

The synthesis of **YD54** and **YDR1** are shown in Scheme 1 and 2, respectively.

**Scheme 1:**
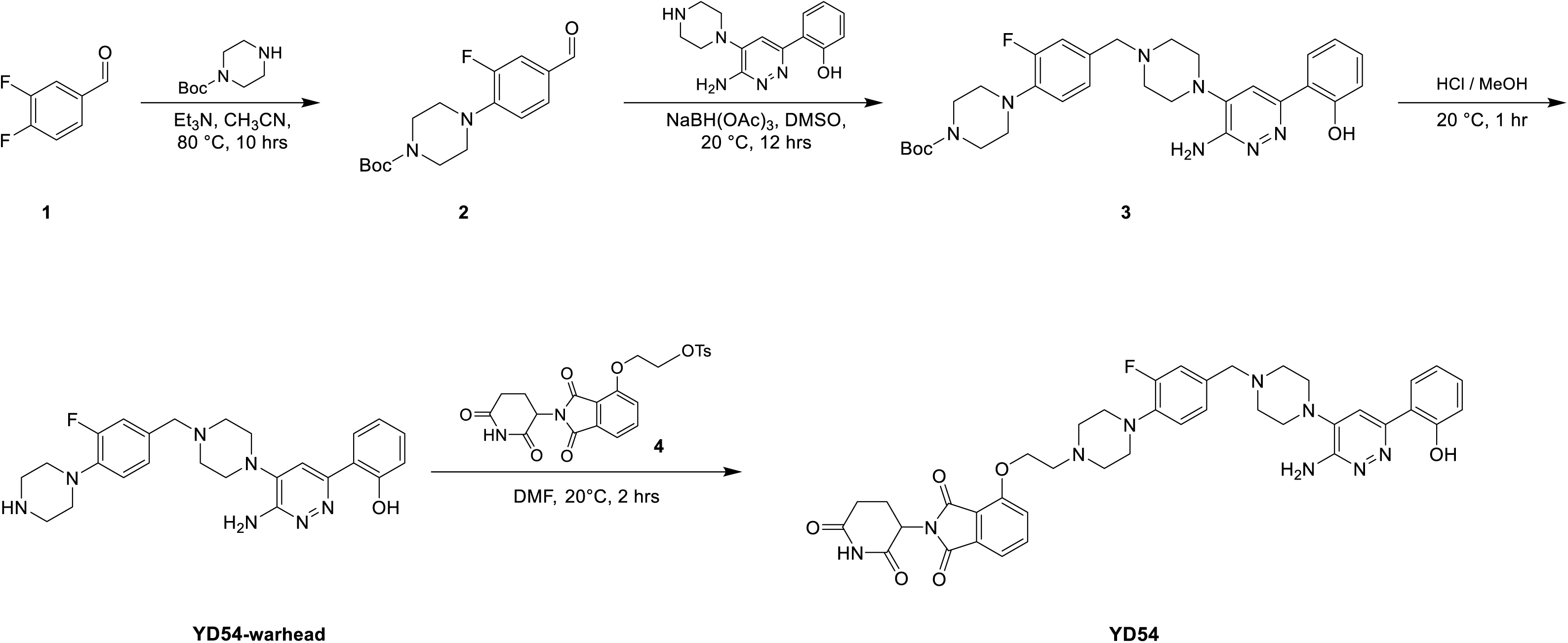
Synthesis of YD54

**Scheme 2:**
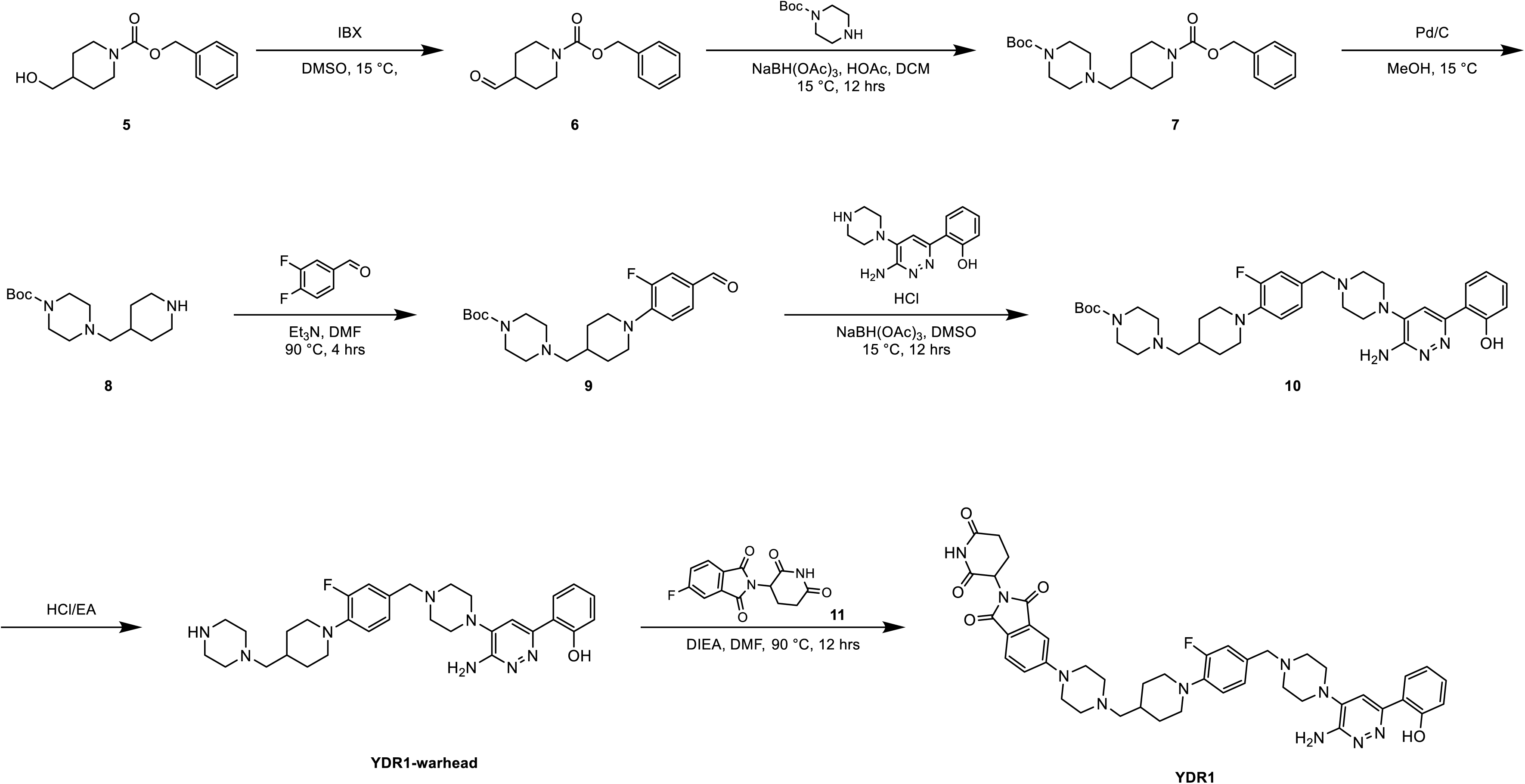
Synthesis of YDR1

**Synthesis of YD54:** Intermediate **2** was prepared from 3,4-difluorobenzaldehyde and Boc-protected piperazine using a SNAr reaction. Following reductive amination with 2-(6-amino-5-(piperazin-1-yl)pyridazin-3-yl) phenol resulted in intermediate **3**. **YD54 warhead** was subsequently obtained by Boc-deprotection in intermediate **3**. The E3 ligase ligand, intermediate **4**, was purchased from LabNetwork. **YD54** was obtained by SNAr reaction between **YD54-warhead** and intermediate **4**.

**Synthesis of YDR1:** Intermediate **6** was obtained by oxidation of alcohol in intermediate **5** to aldehyde, followed by reductive amination with Boc-protected piperazine to make intermediate **7**. The -Cbz group in intermediate **7** was deprotected using hydrogenation to offer intermediate **8**, followed by SNAr reaction with 3,4-difluorobenzaldehyde to prepare intermediate **9**. Another reductive amination was performed between intermediate **9** and 2-(6-amino-5-(piperazin-1-yl)pyridazin-3-yl) phenol to generate intermediate **10**. **YDR1-warhead** was obtained via Boc-deprotection of intermediate **10**. The E3 ligase ligand intermediate **11** was purchased from LabNetwork, and the **YDR1** was obtained by SNAr reaction between **YDR1-warhead** and intermediate **11**.

**Biological activity.** We have performed a comprehensive investigation of the biochemical and cellular effects of YDR1 and YD54 on *SMARCA4* WT and mutant lung cancer cell lines. Initially, we treated H1792, a *SMARCA4*-WT human lung cancer cell line with 1nM to 10µM of YDR1 and YD54 for 24 and 48 hours. YDR1 degraded SMARCA2 in H1792 cells with a half maximal degradation concentration (DC_50_) of 69nM at 24 hours and 60nM at 48 hours while achieving a maximal degradation (D_max_) of 87% at 24 hours and 94% at 48 hours (Figure 2a). YD54 degraded SMARCA2 in H1792 cells with DC**_50_** of 8.1nM at 24 hours and 16nM at 48 hours. YD54 had D_max_ of 98.89% at 24 hours and 99.20% at 48 hours. (Figure 2b). Importantly, YDR1 has only moderate impact on SMARCA4 at 24 hours with DC**_50_** of 135nM and D_max_ of 79% and at 48 hours with DC**_50_** of 381nM and D_max_ of 69% (Figure 2a). YD54 has more impact on SMARCA4 at 24 hours but becomes moderate at 48 hours with DC**_50_** of 149nM and D_max_ of 99.31% (Figure 2b). To further expand the investigation on the potency of YDR1 and YD54 on additional *SMARCA4* mutant cell lines, we treated H322, HCC515, H2030 and H2126 cell lines, with YDR1 and YD54. YDR1 potently degraded SMARCA2 in H322, HCC515, H2030 and H2126 cell lines with a half maximal degradation concentration (DC_50_) of 6.4nM, 10.6nM, 12.7nM and 1.2nM respectively while achieving profound maximal degradation (D_max_) of 99.2%, 99.4%, 98.7% and 99.6% (Figure 2c). Similarly, YD54 potently degraded SMARCA2 in H322, HCC515, H2030 and H2126 cell lines with a half maximal degradation concentration (DC_50_) of 1nM, 1.2nM, 10.3nM and 1.6nM respectively while achieving a maximal degradation (D_max_) of 99.3%, 98.9%, 98.6% and 98.9% (Figure 2d). Taken together, these data show that YDR1 and YD54 are highly potent SMARCA2 degraders.

**Figure 2:**
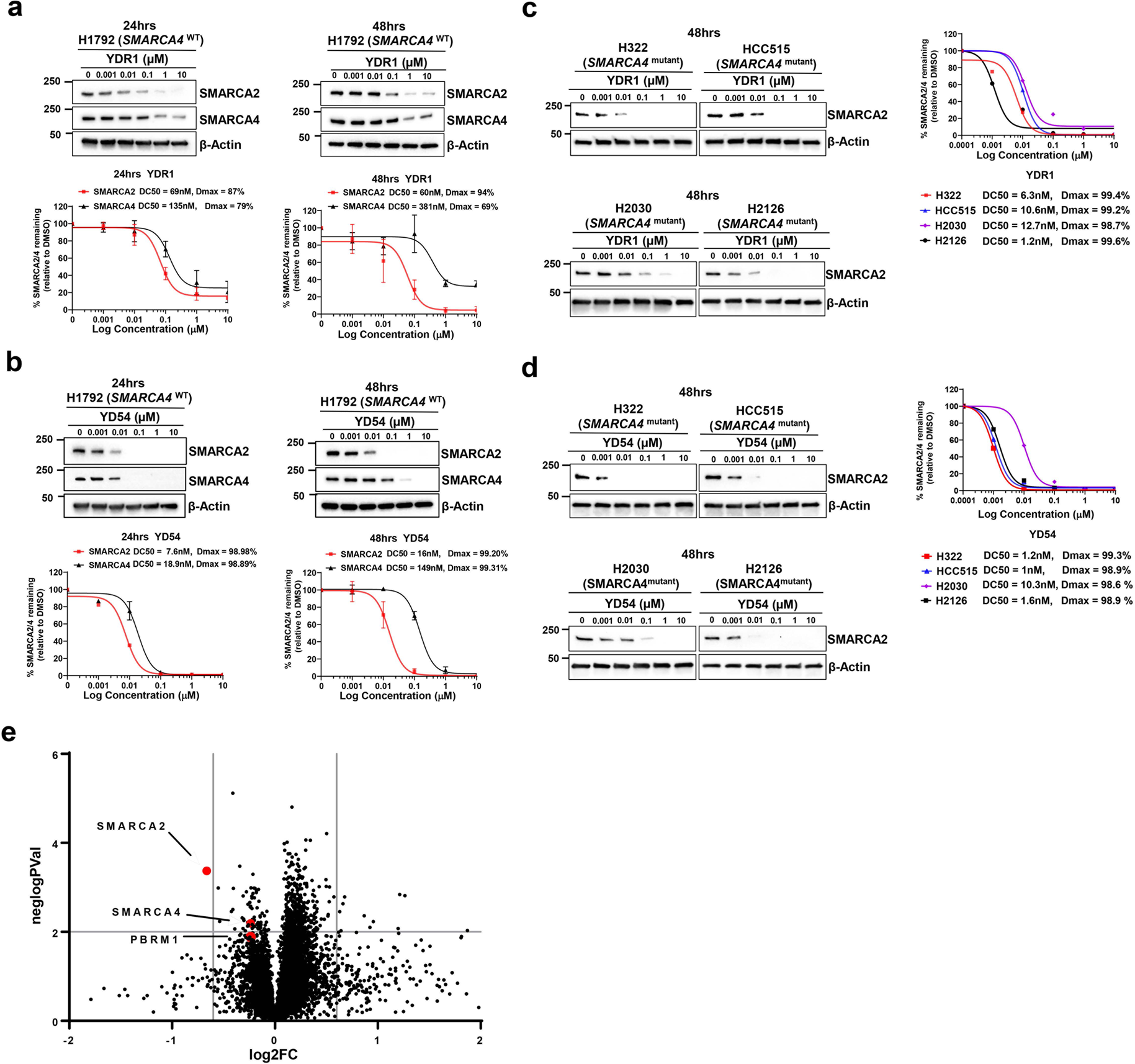
YDR1 and YD54 are potent degraders of SMARCA2. **a,** Immunoblot analysis of SMARCA2 and SMARCA4 in H1792 *SMARCA4*-WT cells following 24hr and 48hr treatments with increasing concentrations of YDR1, as indicated. β-actin serves as a loading control. Quantitation of SMARCA2 and SMARCA4 protein levels by immunoblot analysis in H1792 *SMARCA4*-WT cell line following 24hr and 48hr treatments with increasing concentrations of YDR1, as indicated. Individual DC_50_ values for H1792 cell line were determined from *n* = 3 biologic replicates. Data are normalized to DMSO control-treated cells and presented as mean ± SEM. **b**, Immunoblot analysis of SMARCA2 and SMARCA4 in H1792 *SMARCA4*-WT cells following 24hr and 48hr treatments with increasing concentrations of YD54, as indicated. β-actin serves as a loading control. Quantitation of SMARCA2 and SMARCA4 protein levels by immunoblot analysis in H1792 *SMARCA4*-WT cell lines following 24hr and 48hr treatments with increasing concentrations of YD54, as indicated. Individual DC_50_ values for H1792 cell line were determined from *n* = 3 biologic replicates. Data are normalized to DMSO control-treated cells and presented as mean ± SEM. **c**, Immunoblot analysis of SMARCA2 in H322, HCC515, H2030 and H2126 *SMARCA4*-mutant cell lines following 48hr treatment with increasing concentrations of YDR1, as indicated. β-actin serves as a loading control. Quantitation of SMARCA2 protein levels by immunoblot analysis in H322, HCC515, H2030 and H2126 *SMARCA4*-mutant cell lines following 48hr treatments with increasing concentrations of YDR1, as indicated. **d**, Immunoblot analysis of SMARCA2 in H322, HCC515, H2030 and H2126 *SMARCA4*-mutant cell lines following 48hr treatment with increasing concentrations of YD54, as indicated. β-actin serves as a loading control. Quantitation of SMARCA2 protein levels by immunoblot analysis in H322, HCC515, H2030 and H2126 *SMARCA4*-mutant cell lines following 48hr treatments with increasing concentrations of YD54, as indicated. **e**, Scatterplots of quantitative TMT mass spectrometry of global proteome following 48 hr treatment with 100nM YDR1 in H1792 *SMARCA4*-WT cell lines as indicated. Data are presented as a log2 fold change in the abundance of the respective proteins. Red indicates downregulated SWI/SNF complex related proteins and black indicates downregulated non-SWI/SNF related proteins. A total of 8626 proteins were quantified.

Quantitative proteomics using tandem mass tag (TMT) has rapidly become a powerful approach to investigate the effect of degraders on the cellular proteome to identify targets and establish selectivity profile. Hence, we treated H1792 cell lines with DMSO and 100nM YDR1 for 24 hours and performed TMT mass spectrometry. A total of 8626 proteins were reliably identified and quantified. Fold change and significance plot showed SMARCA2 was the only protein downregulated in YDR1 treated samples at a 1.5-fold change with a *p*-vaule <0.05 showing remarkable selectivity of the compound (Figure 2e). Importantly, SMARCA4 remained minimally altered at these conditions, again highlighting relative selectivity of YDR1 to SMARCA2 over SMARCA4.

### YDR1 and YD54 selectively inhibit growth of *SMARCA4*-mutant lung cancer cell lines

We next assessed the impact of YDR1 and YD54 on cellular growth by performing a 10-day clonogenic assay of two *SMARCA4*-mutant cell lines (H1568 and H1693) and two *SMARCA4*-WT lung cancer cell lines (HCC44 and H2122). Both YDR1 and YD54 caused a dose-dependent inhibition in clonogenic assays in *SMARCA4*-mutant cells (Figure 3a, 3c) with average cellular IC_50_ of 97nM and 11nM respectively (Figure 3b, 3d). Importantly, both YDR1 and YD54 had minimal impact on growth of *SMARCA4* WT cell lines with average cellular IC_50_ of 10µM and 9.1µM respectively (Figure 3b, 3d). This implies about 103-and 827-fold activity of YDR1 and YD54 on *SMARCA4*-mutant cells compared to WT cells showing the remarkable selectivity of cellular phenotypic effect on mutants.

**Figure 3:**
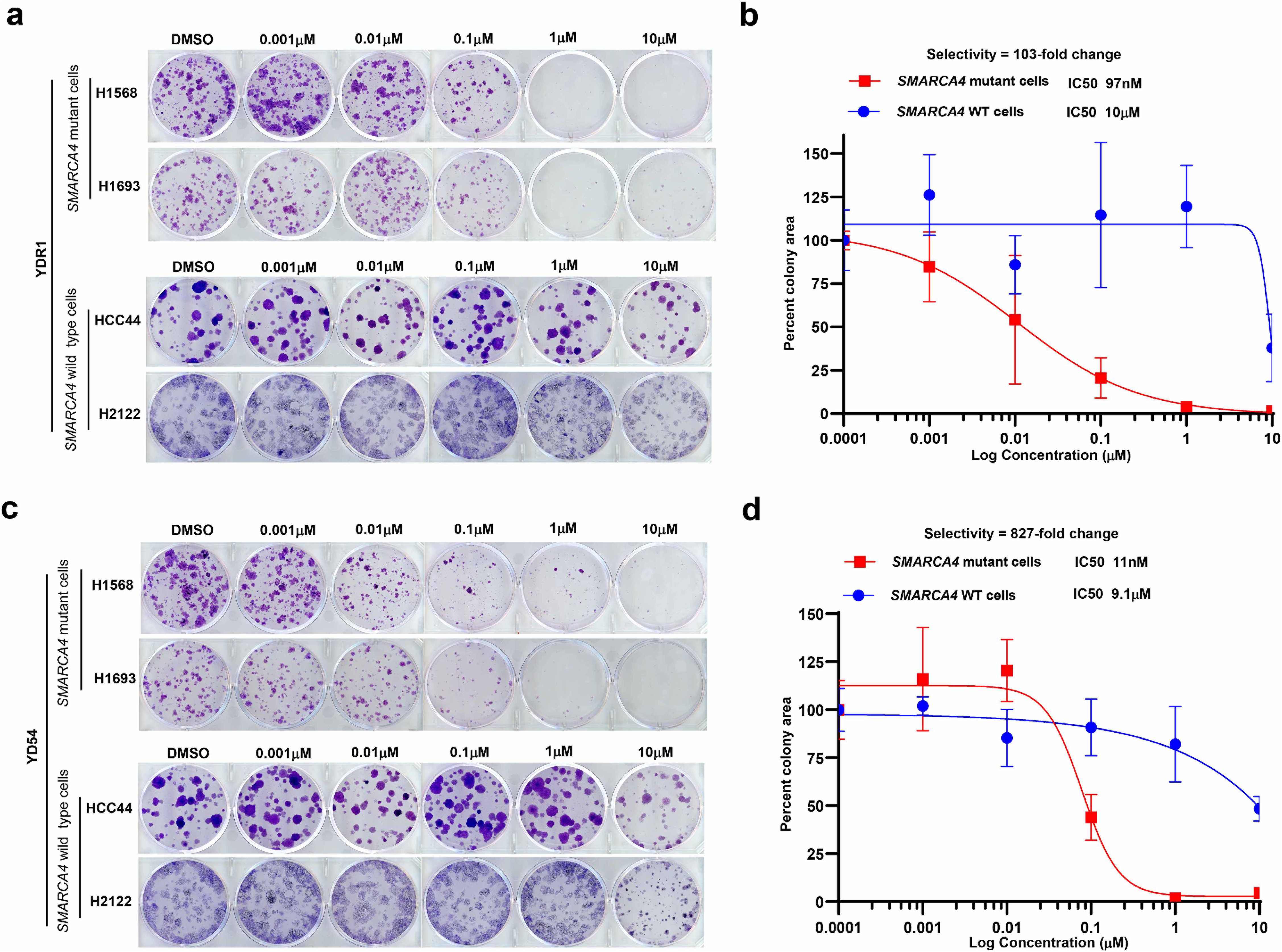
Selective growth inhibitory activity of YDR1 and YD54 in lung cancer cell lines. **a,** Representative images of clonogenic growth assays in H1568 and H1693, *SMARCA4*-mutant cells and HCC44 and H2122, *SMARCA4*-WT cells following 10-12 days of treatment with increasing concentrations of YDR1, as indicated. **b,** Quantitation of percent change in clonogenic area (relative to DMSO) in two *SMARCA4*-mutant and two *SMARCA4*-WT lung cancer cell lines treated with increasing concentrations of YDR1. Error bars represent mean ± SEM from *n* = 3 biologic replicates. **c,** Representative images of clonogenic growth assays in H1568 and H1693, *SMARCA4*-mutant cells and HCC44 and H2122, *SMARCA4*-WT cells following 10-12 days of treatment with increasing concentrations of YD54, as indicated. **d,** Quantitation of percent change in clonogenic area (relative to DMSO) in two *SMARCA4*-mutant and two *SMARCA4*-WT lung cancer cell lines treated with increasing concentrations of YD54. Error bars represent mean ± SEM from *n* = 3 biologic replicates.

### Metabolic stability and pathways of inactivation of YDR1 and YD54 in liver microsomal assays

For efficient activity, PROTACs need to induce proximity of target protein and E3 ligase. An excess of E3 ligase binding ligand or protein of interest ligand can effectively compete for formation of ternary complex. A recent report highlighted metabolites of PROTACs can potentially compete for binding to ligand or E3 ligase prevent full PROTAC mediated degradation^32^. Hence, we were interested to explore the stability and possible metabolites of YDR1 and YD54 and performed human and mouse liver microsomal stability assays followed by metabolite identification. YDR1 shows slow clearance with T_1/2_ of 58 minutes while YD54 shows moderate clearance with T_1/2_ of 35 minutes in human microsomal assays (Table 1). Similar trends were observed in mouse microsomal assays with YDR1 having T_1/2_ of 59 minutes and YD54 having 46 minutes (Table 1). We next sought to identify and characterize major metabolites of YDR1 and YD54 in mouse liver microsomes. Compounds at 10µM were incubated with mouse liver microsomes at 37°C for 60 minutes. After incubation, samples were analyzed by LC-UV-HRMS and the structures of the metabolites were proposed based on the interpretation of their MS and MS^2^ data. Under these experimental conditions, 3 metabolites were tentatively identified for YDR1 in mouse liver microsomes (Figure 4a). M1: Mono-oxygenation and hydrogenation metabolite (P + O + 2H); M2: Mono-oxygenation metabolite (P + O); M3: Oxidative dealkylation and mono-oxygenation metabolite (P + O – C_14_H_17_N_5_O + O). In mouse liver microsomes, M2 was considered to be the top one metabolite with relative abundance of 1.02%. Similarly, under the current experimental conditions, 3 metabolites were tentatively identified for YD54 in mouse liver microsomes. M1: *N*-Dealkylation metabolite (P – C_15_H_12_N_2_O_5_); M2: Mono-oxygenation and *N*-dealkylation metabolite (P + O – C_15_H_12_N_2_O_5_); M3: Hydrolysis metabolite (P + H_2_O). In mouse liver microsomes, M1 was considered to be the top one metabolite with relative abundance of 9.24% (Figure 4b). These data suggest that minimal metabolite-competition is expected in YDR1 while modest competition is expected from the dealkylation metabolite M1 in the case of YD54.

**Table 1.**
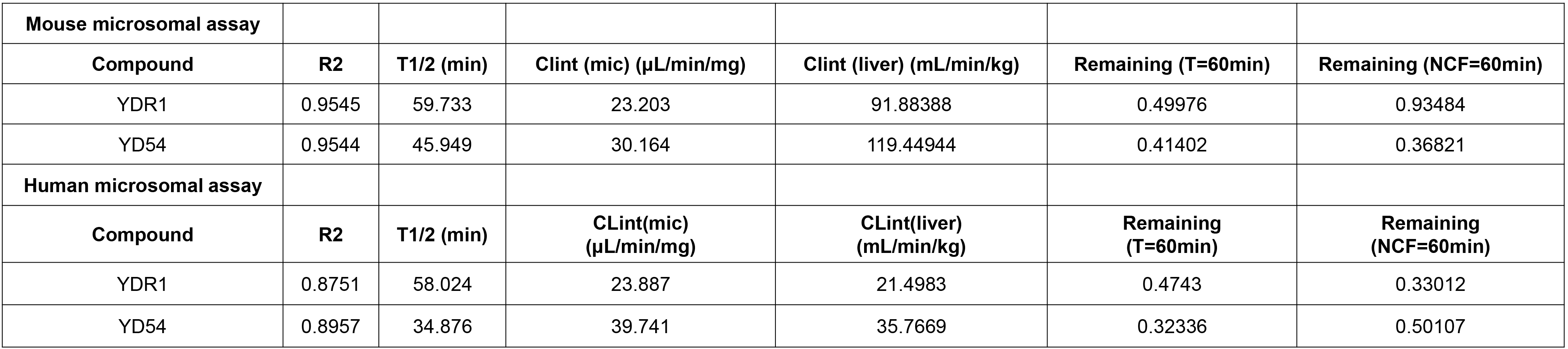
Design of the mesocomic experiments.

**Table 2.**

|  |  |  |  |  |
| --- | --- | --- | --- | --- |
|  | YDR1 P.O | YDR1 IP | YD54 P.O | YD54 IP |
| PK Parameters | Mean | Mean | Mean | Mean |
| C<sub>max</sub> (ng/mL) | 649.3 | 1756.3 | 587.0 | 2039.7 |
| T<sub>1/2</sub> (h) | ND | 44.8 | 6.0 | 13.0 |
| AUC<sub>0-last</sub> (ng.h/mL) | 12079.3 | 23213.7 | 4743.0 | 19365.3 |
| AUC<sub>0-inf</sub> (ng.h/mL) | ND | 68813.0 | 5122.0 | 27108.3 |

**Figure 4:**
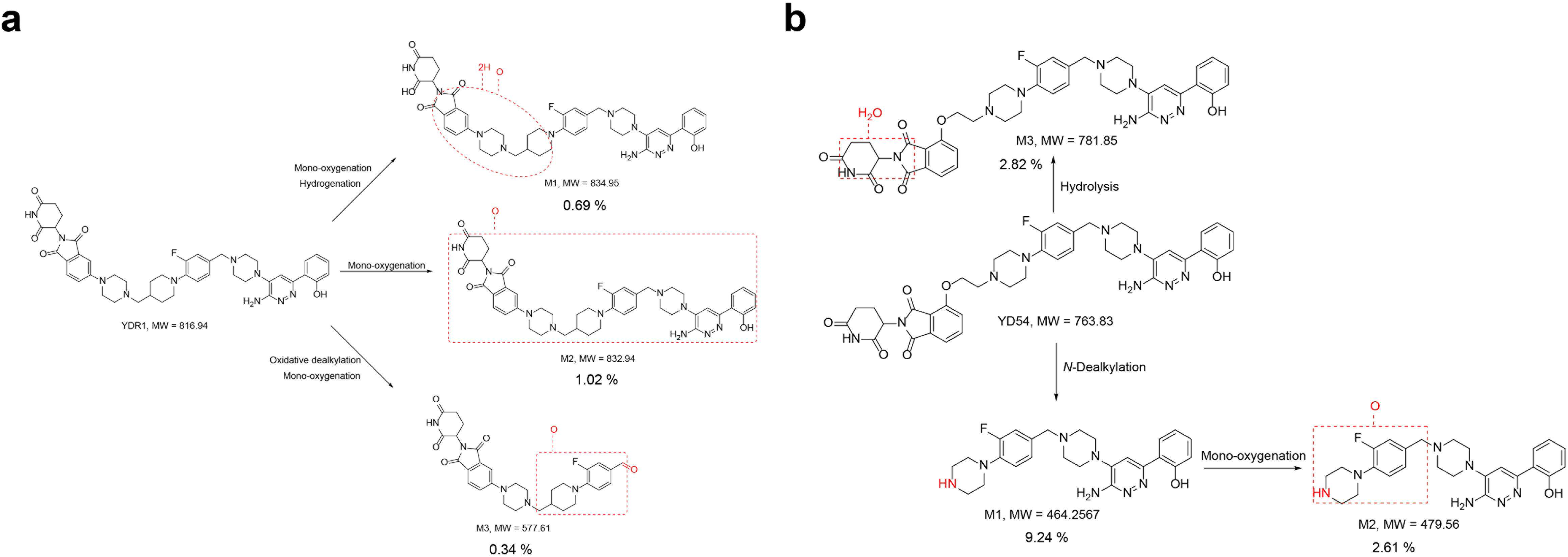
Metabolic stability and pathways of inactivation of YDR1 and YD54 in liver microsomal assays. **a**, 10µM concentration of YDR1 and YD54 were incubated with mouse liver microsomes at 37°C for 60 minutes and samples was analyzed by LC-UV-HRMS and percentage of each metabolite structures were indicated. **b,** YD54 of concentration 10µM was incubated with mouse liver microsomes at 37°C for 60 minutes, samples were analyzed by LC-UV-HRMS and percentage of each metabolite structures were indicated.

### YDR1 and YD54 are orally bioavailable

We next determined the bioavailability of YDR1 and YD54 after intraperitoneal (IP) or oral routes of administration. CD-1 mice were dosed with 25mg/kg of YDR1 or YD54 in 0.5% methylcellulose/0.2%Tween 80 in water (for oral) or in 5%DMSO/5% solutol/90% water (for IP). Plasma was collected at 0.25, 0.5, 1, 2, 4, 8 and 24 hours and amount of YDR1 and YD54 determined by LC-MS/MS. Pharmacokinetic parameters are tabulated in Table 2. We observed robust oral bioavailability and stability of YDR1 with T_1/2_ value exceeding 24 hours (Table 2 and Figure 5). YD54 was also orally bioavailable with faster clearance as predicted from liver microsomal studies with T_1/2_ of 6 hours (Table 2 and Figure 5).

**Figure 5:**
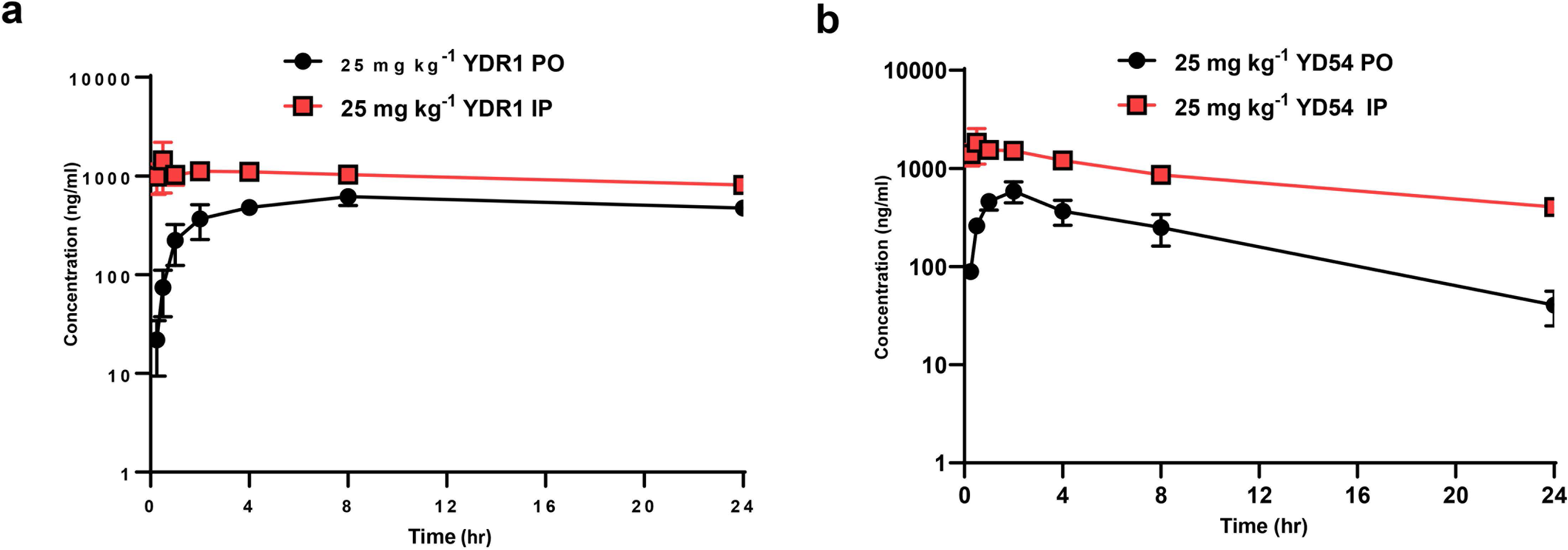
YDR1 and YD54 are orally bioavailable. **a,** Plasma concentrations of YDR1 (25 mg kg^-1^) in male CD-1(ICR) mice following a single p.o and i.p dose over 24 hours. Plasma was collected at 0.25, 0.5, 1, 2, 4, 8 and 24 hours and amount of YDR1 and YD54 determined by LC-MS/MS. **b,** Plasma concentrations of YD54 (25 mg kg^-1^) in mice following a single p.o dose and i.p dose over 24 hours. Plasma was collected at 0.25, 0.5, 1, 2, 4, 8 and 24 hours and amount of YD54 determined by LC-MS/MS. Values are shown as mean ± SEM (n =3 mice for each time point).

### Pharmacokinetic (PK) and pharmacodynamic (PD) correlation studies of YDR1

After establishing basic PK profiles, we performed in depth PK/PD correlation studies using western blot quantification of *in vivo* levels of SMARCA2 as key target engagement parameter. First, we administered vehicle control, 5mg/kg, 10mg/kg, 20mg/kg, 40mg/kg or 80mg/kg YDR1 orally, once daily for 3 days in C57BL/6 mice. 24 hours after last administration, mice were euthanized, plasma concentration of YDR1 measured and spleen tissue was analyzed by western blot for SMARCA2 expression. We observed dose dependent increase in plasma concentration of YDR1 ranging from about 100nM in the 5mg/kg group to 1.88µM in the 80mg/kg group (Figure 6a). Importantly, dose dependent degradation of SMARCA2 with highest degradation of 70% noted in the 80mg/kg level (Figure 6b). While this was encouraging, it is below the level of degradation observed in human cancer cells *in vitro*. It is important to note that we have utilized pomalidomide as E3 ligand recruiter in our PROTACs and it is known that in mice thalidomide and its analogs generally have lower affinity to mouse Cereblon due to the presence of isoleucine at position 391 in contrast to valine in humans. This has been demonstrated to result in steric clash and reduced affinity to mouse Cereblon. This has inspired the development of genetically engineered mice harboring I391V substitution (*Crbn ^I^*^391^*^V^*) resulting in better recapitulation of effect of thalidomide and analogs on human cells and tissues ^33, 34^. Hence, we next assessed the impact of our Cereblon-dependent PROTAC YDR1 in *Crbn ^I^*^391^*^V^* mice. Specifically, we administered vehicle control, 10mg/kg, 20mg/kg, 40mg/kg or 80mg/kg YDR1 orally, once daily for 3 days in *Crbn ^I^*^391^*^V^* mice. 24 hours after last administration, mice were euthanized, plasma concentration of YDR1 measured and spleen tissue was analyzed by western blot for SMARCA2 expression. We observed dose dependent increase in plasma concentration of YDR1 ranging from 1µM in the 10mg/kg group to 2.45µM in the 80mg/kg group (Figure 6c). Importantly, more robust dose dependent degradation of SMARCA2 with highest degradation of 81% was noted in the 80mg/kg dose, (Figure 6d). At all doses tested, we see increased degradation of SMARCA2 (range of 11% to 38%) in *Crbn ^I^*^391^*^V^* mice compared to C57BL/6 mice, indicating the usefulness of *Crbn ^I^*^391^*^V^* mice for more accurate modeling of effect of Cereblon-based PROTACs.

**Figure 6:**
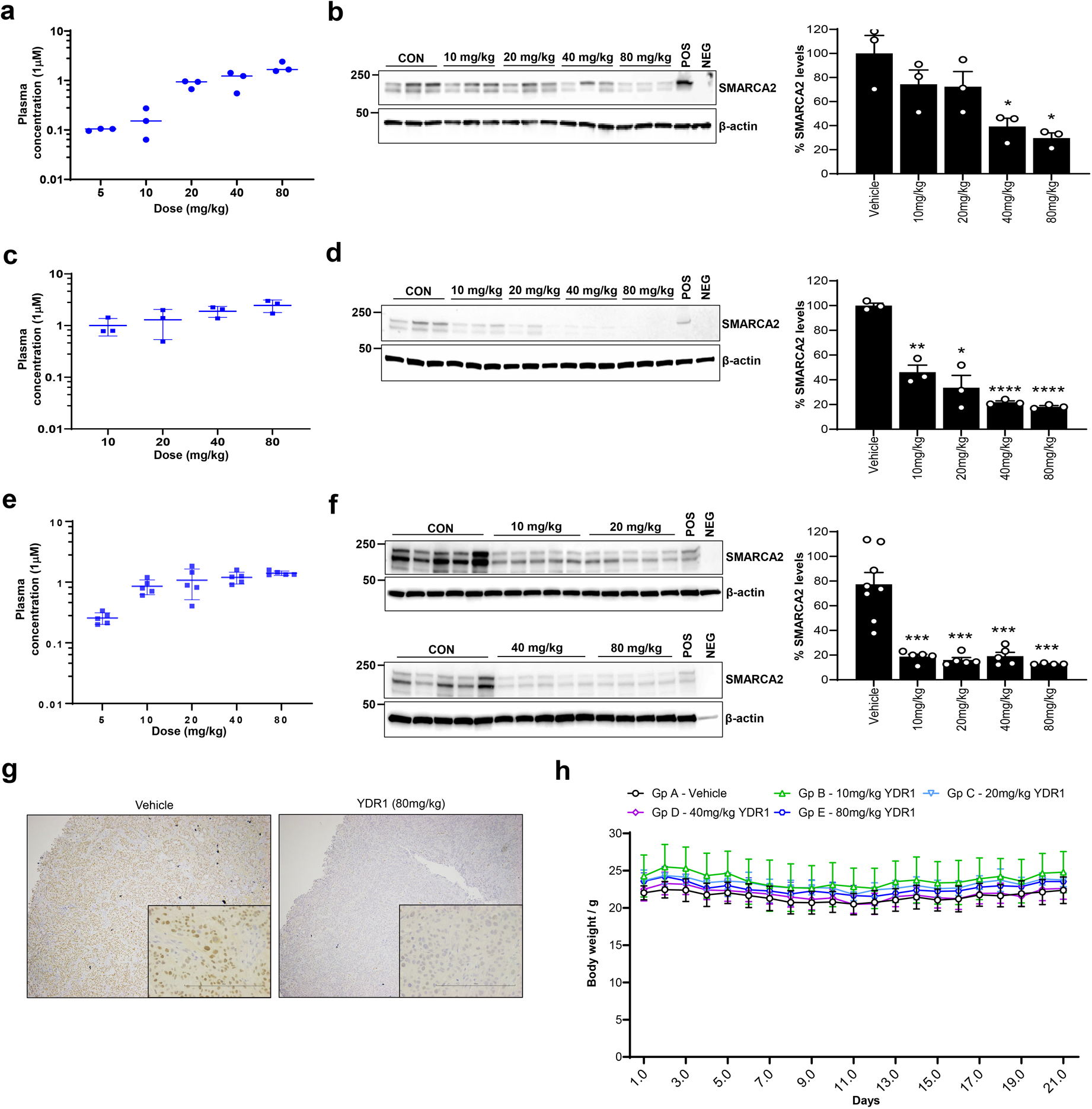
Pharmacokinetic and pharmacodynamic correlation studies of YDR1 and YD54. **a)** Plasma concentrations of YDR1 (5 mg kg^-1^, 10 mg kg^-1^, 20 mg kg^-1^ 40 mg kg^-1^ and 80 mg kg^-1^) in C57BL/6 mice following multiple p.o dose over 24 hours. Values are shown as mean ± SEM (n =3 mice for each time point). **b**) Dose-dependent degradation of SMARCA2 by YDR1 in spleen of C57BL6 mice treated with YDR1 (5 mg kg^-1^ 10 mg kg^-1^ 20 mg kg^-1^ 40 mg kg^-1^ and 80 mg kg^-1^ . Quantification of SMARCA2 protein levels normalized to loading control β-actin, from immunoblots in (**b**) was done by ImageJ software. Values are shown as mean ± SEM (n =5 mice for each group). *P* values were calculated from two-tailed unpaired Student’s t-tests between vehicle and treatment group. **c)** Plasma concentrations of YDR1 (5 mg kg^-1^ 10 mg kg^-1^ 20 mg kg^-1^, 40 mg kg^-1^ and 80 mg kg^-1^) in humanized *Crbn ^I^*^391^*^V^* mice following multiple p.o dose over 24 hours. Values are shown as mean ± SEM (n =3 mice for each time point). **d**) Dose-dependent degradation of SMARCA2 by YDR1 in Spleen of *Crbn ^I^*^391^*^V^* mice treated with YDR1 (10 mg kg^-1^, 20 mg kg^-1^, 40 mg kg^-1^ and 80 mg kg^-1^ . Quantification of SMARCA2 protein levels normalized to loading control β-actin, from immunoblots in (**d**) was done by ImageJ software. Values are shown as mean ± SEM (n =5 mice for each group). *P* values were calculated from two-tailed unpaired Student’s t-tests between vehicle and treatment group). **e)** Plasma concentrations of YDR1 (10 mg kg^-1^, 20 mg kg^-1^, 40 mg kg^-1^ and 80 mg kg^-1^) in NU/J nude (H1568 xenograft) mice following multiple p.o dose over 24 hours. Values are shown as mean ± SEM (n =3 mice for each time point). **f**) Dose-dependent degradation of SMARCA2 by YDR1 in tumor of NU/J nude mice (H1568 xenograft) treated with YDR1 (5 mg kg^-1^, 10 mg kg^-1^, 20 mg kg^-1^, 40 mg kg^-1^ and 80 mg kg^-1^ _)_. Quantification of SMARCA2 protein levels normalized to loading control β-actin, from immunoblots in (**f**) was done by ImageJ software. Values are shown as mean ± SEM (n =5 mice for each group). *P* values were calculated from two-tailed unpaired Student’s t-tests between vehicle and treatment group. (**g)** Tumor IHC images of SMARCA2 staining in H1568 xenografts from vehicle and YDR1 (80 mg kg^-1^) treated mice collected after 24hours at end of the study from mice in (**e**). Scale bar in inset is 200μm. (**h**) Body weight changes of nude mice following daily p.o administration of YDR1 ( 10 mg kg^-1^, 20 mg kg^-1^, 40 mg kg^-1^ and 80 mg kg^-1^) or vehicle-treated control for indicated 21 days. The vehicle is 0.5% Methylcelluose + 1% Tween-80. Data is presented as mean ± SEM.

Next, we sought to determine the ability of YDR1 to induce degradation of SMARCA2 in xenograft tumors. We implanted H1568 human lung cancer cells in the flanks of immunocompromised mice and performed YDR1 administration daily at 10mg/kg, 20mg/kg, 40mg/kg and 80mg/kg for 4 days daily by oral gavage. Tumors were harvested and SMARCA2 probed by western blot. YDR1 potently degraded SMARCA2 in this model with 87% degradation observed at 80mg/kg (Figure 6 e-f). We further validated this by robust reduction in SMARCA2 signal by immunohistochemical staining of tissue sections with SMARCA2 antibodies (Figure 6g). Lastly, we determined the medium-term tolerability of YDR1 by daily treatment of mice with increasing doses up to 80mg/kg for 21 days. Overall, YDR1 was well tolerated in these mice including at the highest dose, with minimal body weight loss and no obvious signs of toxicity or morbidity (Figure 6h).

### Anti-tumor activity of YDR1 in *SMARCA4*-mutant models of NSCLC

Having thus established the basic pharmacokinetics, tolerability and target engagement of YDR1, we finally sought to determine its anti-tumor inhibitory activity using human xenograft tumor models. We first tested YDR1 in H1568 xenograft model (*SMARCA4*-mutant) in female athymic nude and co-administered 40mg/kg YDR1 once daily by oral gavage for 14 days. We found that YDR1 at 40m/kg moderately inhibited tumor growth with a tumor growth inhibition (TGI) of 25.2 % compared to vehicle control (Figure 7a). We also tested YD54 in H2023 xenograft model (*SMARCA4*-mutant) in female athymic nude and co-administered 5mg/kg YD54 once daily by oral gavage for 19 days. We found that YD54 at 5mg/kg inhibited tumor growth with a tumor growth inhibition (TGI) of 54.7% compared to vehicle control (Figure 7d). In both the anti-tumor efficacy experiments, YDR1 and YD54 caused no or minimal reduction in body weight (<10% compared to controls) and were tolerated at the doses and duration of the *in vivo* studies (Figure 7b, e). Immunoblot analysis of individual tumor protein lysates from this study showed YDR1 and YD54 reduced SMARCA2 levels by 50% and 76% respectively when compared to vehicle control-treated tumors (Figure 7c,7f).

**Figure 7:**
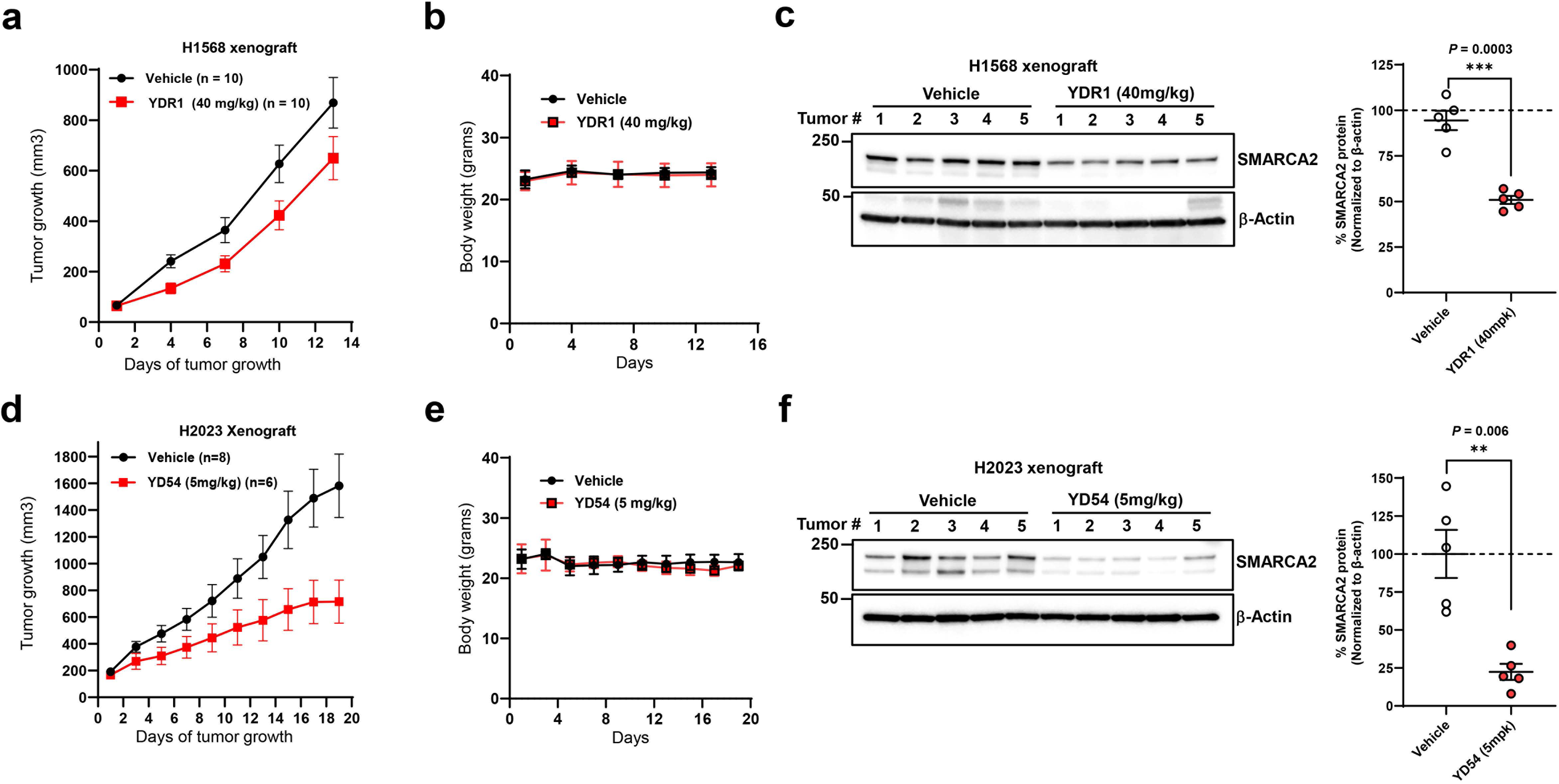
Anti-tumor efficacy of YD54 in *SMARCA4*-mutant xenograft models of lung cancer. **a**, Anti-tumor efficacy of nude mice harboring H1568 *SMARCA4*-mutant xenografts following daily administration of YDR1 (40 mg kg^-1^) or vehicle treated control for 14 days. YD54 and vehicle was administered by p.o route. The vehicle is 0.5% methylcelluose + 1% Tween-80. Data is presented as mean ± SEM. **b**) Body weight changes of nude mice following daily p.o administration of YDR1 (40 mg kg^-1^) or vehicle-treated control. Data is presented as mean ± SEM. **c)** Dose-dependent degradation of SMARCA2 by YD54 in tumor of nude mice (H1568 xenograft) treated with YDR1 (40 mg kg^-1^). Quantification of SMARCA2 protein levels normalized to loading control β-actin, from immunoblots in (**c**) was done by ImageJ software. Values are shown as mean ± SEM (n =5 mice for each group). *P* values were calculated from two-tailed unpaired Student’s t-tests between vehicle and treatment group. **d**, Anti-tumor efficacy of nude mice harboring H2023 *SMARCA4*-mutant xenografts following daily administration of YD54 (5 mg kg^-1^) or vehicle treated control for 19 days. YD54 and vehicle was administered by p.o route. The vehicle is 0.5% methylcelluose + 1% Tween-80. Data is presented as mean ± SEM. **e**) Body weight changes of nude mice following daily p.o administration of YD54 (5 mg kg^-1^) or vehicle-treated control. Data is presented as mean ± SEM. **f)** Dose-dependent degradation of SMARCA2 by YD54 in tumor of nude mice (H2023 xenograft) treated with YD54 ( 5 mg kg^-1^). Quantification of SMARCA2 protein levels normalized to loading control β-actin, from immunoblots in (**f**) was done by ImageJ software. Values are shown as mean ± SEM (n =5 mice for each group). *P* values were calculated from two-tailed unpaired Student’s t-tests between vehicle and treatment group.

### YDR1 and YD54 synergizes with sotorasib to inhibit growth of sotorasib-resistant *KRAS^G12C^* **and *SMARCA4* co-mutant cancer cells.**

It is worth noting that YDR1 did not induce tumor regression in any of our experiments which is consistent with other anti-tumor efficacy studies employing other SMARCA2 PROTACs^23, 24^ , suggesting identification of synergistic combinations are required for efficient clinical translation. Importantly, a recent clinical report showed that co-mutations in tumor suppressors (*SMARCA4, KEAP1 and CDKN2A*) predict poorer response to KRAS^G12C^ inhibitors in lung cancer patients ^35^. This study identified both inferior overall survival (OS) and progression-free survival (PFS) among patients with *SMARCA4* mutation (PFS: 1.6 vs 5.4 months, OS: 4.9 vs. 11.8 months, for mutant vs WT *SMARCA4*). Interestingly, an elegant study of *KRAS* mutant rhabdoid tumors with concomitant loss of *SMARCB1* (*another key SWI/SNF subunit*) showed lack of activation of MAPK signaling pathway indicating SWI/SNF inactivation can lead to “partial independence” from upstream RAS pathway^36^. We initially corroborated the clinical data that suggested that *SMARCA4* mutant tumors have poor response to KRAS^G12C^ inhibitors by analyzing the vast functional genomics data from the DepMap Project^37^ that assessed the dependency of lung cancer cell lines for growth and survival on *KRAS*. We categorized *KRAS* mutant lung cancer cell lines by *SMARCA4* status and asked if there is a difference in genetic dependency on *KRAS*. We showed that indeed *SMARCA4* and *KRAS^G12C^* co-mutant cell lines are significantly less dependent on *KRAS* than *SMARCA4* WT cells (Figure 8a). Hence, we hypothesized that co-treatment of tumors harboring both *KRAS^G12C^* and *SMARCA4* mutations with the combination of SMARCA2 PROTAC and KRAS^G12C^ inhibitor might have synergistic effect. We used H2030 cell line which harbors *KRAS ^G12C^* mutation and *SMARCA4* deletion for our investigations. We treated these cells with sotorasib, YDR1 or combination of sotorasib and YDR1 and performed a 7-day growth assay. While we observed minimal growth inhibition by the single agents, the combination induced a robust synergistic growth inhibition (Figure 8b). We next confirmed the statistical significance of growth inhibition by the combination of YDR1 and sotorasib by estimating the Bliss/Lowe consensus synergy score (>10 indicates synergy) from SynergyFinder^38^ (Figure 8c). We corroborated this by performing 14-day clonogenic assays which similarly showed robust and significant synergy in the combination of sotorasib and YDR1 (Figure 8d-f). We also performed clonogenic assays with H2030 cells treated with sotorasib, YD54 or combination of sotorasib and YD54 to know if the effect is synergistic like YDR1. We confirmed the statistical significance of growth inhibition by the combination of YD54 and sotorasib by estimating the Bliss/Lowe consensus synergy score (>10 indicates synergy). We observed more synergistic effect of combination of YD54 and Sotorasib than combination of YDR1 and Sotorasib. The demonstration of synergistic combination effect of YDR1 or YD54 with sotorasib in a sotorasib-resistant H2030 cell line is highly promising and could enable expansion of the therapeutic indications for both SMARCA2 PROTACs and KRAS^G12C^ inhibitors.

**Figure 8:**
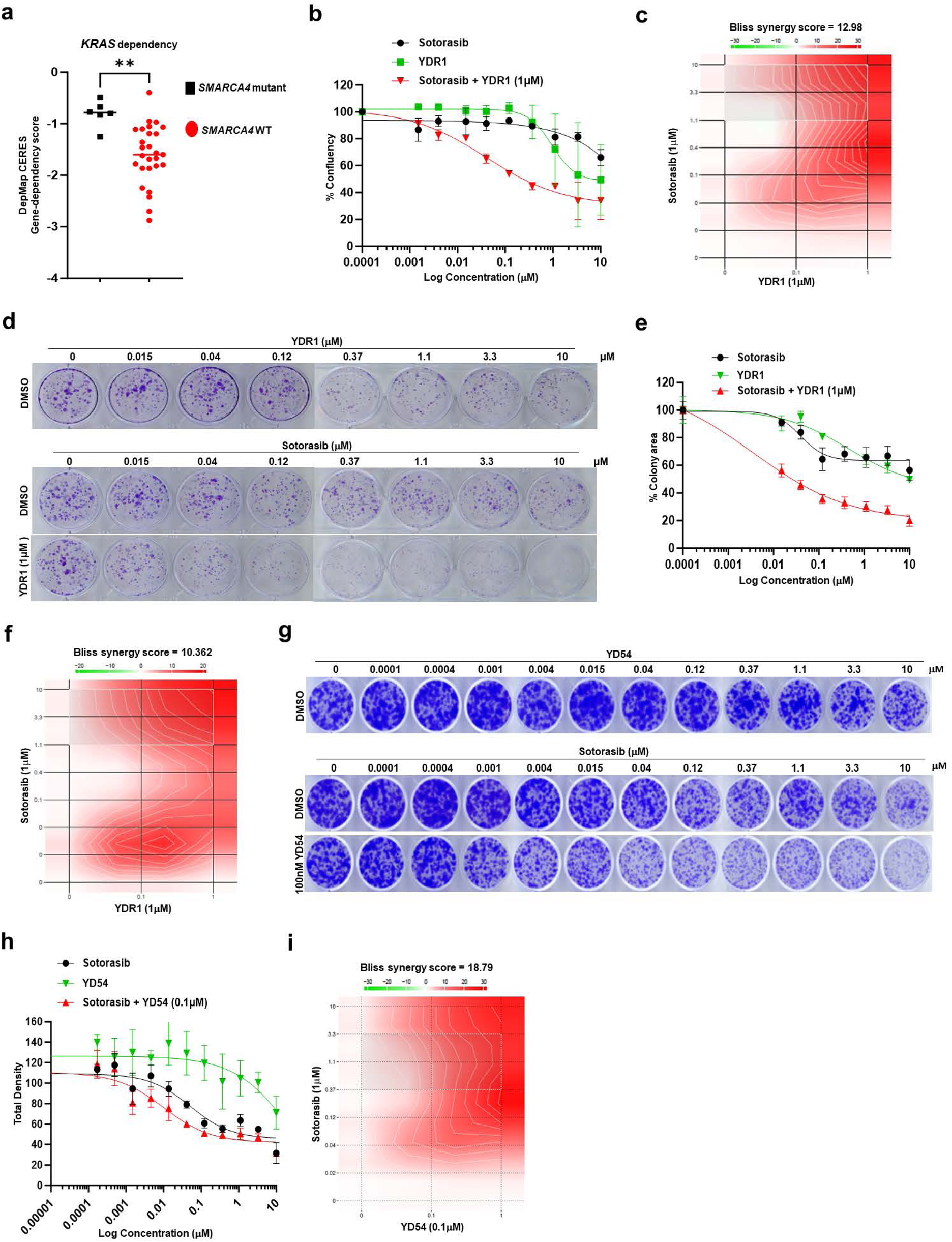
YDR1 and YD54 synergize with sotorasib to inhibit growth of sotorasib-resistant *KRAS^G12C^* and *SMARCA4* co-mutant cancer cells. **a)** Graph shows *KRAS* dependency on *SMARCA4*-WT cell lines or *SMARCA4*-mutant cell lines. **b)** Cell proliferation (% confluency) of H2030 *SMARCA4*-mutant cell line was determined by Incucyte after treatment with either 3-fold diluted concentrations of Sotorasib or YDR1 or combination treatment of 1mM YDR1 and doses of Sotorasib. Error bars represent mean ± SEM from *n* = 3 biologic replicates. **c)** Heatmaps of H2030 cell line depicts synergistic drug interactions using the Bliss independence model. Treatment of H2030 *SMARCA4*-mutant cell line, as indicated (b). A score > 10 is considered as synergy. A score between -10 to 10 is considered additive. A score < -10 is considered antagonistic. **d**) Representative images of clonogenic growth assays in H2030 cells treated with either 3-fold diluted concentrations of Sotorasib or YDR1 or combination treatment of 1μM YDR1 and doses of Sotorasib. Error bars represent mean ± SEM from *n* = 3 biologic replicates. **e)** Quantitation of percent colony area (relative to DMSO) in H2030 *SMARCA4*-mutant lung cancer cell line treated as in (d), Error bars represent mean ± SEM from *n* = 3 biologic replicates. **f)** Heatmaps of H2030 cell line depicts synergistic drug interactions using the Bliss independence model. Treatment of H2030 *SMARCA4*-mutant cell line, as indicated (e). A score > 10 is considered a synergy. A score between -10 to 10 is considered additive. A score < -10 is considered antagonistic. (**g**) Representative images of clonogenic growth assays in H2030 *SMARCA4*-mutant cell line cell line treated with either 3-fold diluted concentrations of Sotorasib or YD54 or combination treatment of 0.1μM YD54 and doses of Sotorasib. Error bars represent mean ± SEM from *n* = 3 biologic replicates. **h)** Quantitation of total density (relative to DMSO) in H2030 *SMARCA4*-mutant lung cancer cell line treated as in (g), Error bars represent mean ± SEM from *n* = 3 biologic replicates. **i)** Heatmaps of H2030 cell line depicts synergistic drug interactions using the Bliss independence model. Treatment of H2030 *SMARCA4*-mutant cell line, as indicated (h). A score > 10 is considered a synergy. A score between -10 to 10 is considered additive. A score < -10 is considered antagonistic.

Taken together, YDR1 and YD54 were identified as novel SMARCA2 PROTACs that selectively degrade SMARCA2 allowing for an efficacious anti-tumor effect as single agent or in combination with KRAS^G12C^ inhibitors in *SMARCA4* mutant cancers.

## DISCUSSION

Targeted protein degradation is a rapidly evolving therapeutic modality with substantial potential in cancer treatment. Human and mouse genetics as well as pharmacologic evidences have firmly established that *SMARCA2* is synthetic lethal to *SMARCA4,* required for growth and survival of *SMARCA4*-mutant cancer cells and a highly desirable therapeutic target ^16, 17, 19, 20, 23,24^. The strength of the genetic interaction coupled to the large number of patients harboring *SMARCA4* mutations across multiple cancer types strongly suggest that therapeutics targeting this synthetic lethality could have paradigm shifting consequence like PARP inhibitors in *BRCA*-mutant tumors. Hence, there is intense effort to develop SMARCA2 inhibitors and take advantage of this vulnerability of *SMARCA4*-mutant cancers. So far, the major challenge has been the fact that the readily ligand accessible SMARCA2 bromodomain is not required for this synthetic lethal interaction^16^, hence mere inhibition of the bromodomain has no therapeutic utility. Since potent SMARCA2 bromodomain ligands exist^29^, PROTACs provide a unique opportunity to utilize these bromodomain ligands as hooks and enable degradation of the entire protein to achieve therapeutic utility. To date, multiple potent SMARCA2 PROTACs have been reported^17, 23–25^. Importantly, these PROTACs have utilized the VHL E3 ubiquitin ligase and shown promising anti-tumor activities. Since one of the major mechanisms of cancer resistance to PROTACs is genomic alterations in the E3 ligase targeted by the PROTAC, it is highly desirable to have a toolbox of PROTACs with non-overlapping E3 ligases^28^.

With the goal of expanding the repertoire of SMARCA2 PROTACs using alternative E3 ligases, we report the discovery of YD54 and YDR1, two potent and selective SMARCA2 degrading PROTACs based on the E3 ligase Cereblon. Optimization of linkers, aptly named linkerology, is a key factor in PROTAC design^39, 40^, and in our studies rigid heterocyclic ring containing linkers were found to improve potency, metabolic stability and pharmacokinetic properties of PROTACs. We performed in depth biochemical, cellular and pharmacologic characterization of these compounds and showed that they have highly selective growth inhibitory activity against *SMARCA4* mutant cancer cells (100 to 800-fold).

Selective SMARCA2 degraders (with limited activity against SMARCA4) are desirable for clinical development due to the anticipated toxicities of unintended SMARCA4 degradation. This is borne out of mouse knockout studies that showed essentiality of SMARCA4 for tissue maintenance as well as overt toxicity in mice by pharmacologic inhibitors of SMARCA4 ATPase activity^21, 41, 42^. YDR1 and YD54 show enhanced selectivity for SMARCA2 over SMARCA4 by immunoblotting. Furthermore, at the global level, quantitative proteomics showed that SMARCA2 is the most significantly degraded protein among 8626 proteins analyzed while SMARCA4 was not. YDR1was also well tolerated in mice up to 21 days of treatment suggesting existence of likely therapeutic window. Importantly, we showed that YDR1 and YD54 were orally bioavailable and show potent SMARCA2 degradation profile in mouse tissues as well as in implanted xenograft tumors. Lastly, we demonstrated robust anti-tumor growth inhibition in a *SMARCA4*-mutant xenograft model of lung cancer.

Finally, we were inspired by a recent clinical observation that showed that co-mutations in tumor suppressors such as *SMARCA4* predict poorer response to KRAS^G12C^ inhibitors in lung cancer patients ^35^ to conduct combination studies. We were able to test and confirm our hypothesis that co-treatment of tumors harboring both *KRAS^G12C^* and *SMARCA4* mutation with the combination of SMARCA2 PROTAC and KRAS^G12C^ inhibitor has synergistic growth inhibitory effect. Our demonstration of synergistic combination effect of YDR1 or YD54 with sotorasib in a sotorasib-resistant H2030 cell line is highly promising and could enable expansion of the therapeutic indications of SMARCA2 PROTACs and KRAS^G12C^ inhibitors.

## CONCLUSIONS

In summary, we discovered a novel series of SMARCA2 PROTACs based on Cereblon E3 ligase harboring rigid heterocyclic ring-based linkers. Quantitative proteomics showed these compounds showed enhanced selectivity for degrading SMARCA2 over SMARCA4. Further, in vitro growth assays showed profound selectivity of YD series SMARCA2 PROTACs for *SMARCA4*-mutant cancer cells. Importantly, YDR1 was orally bioavailable and well-tolerated in mice. Additionally, YDR1 and YD54 were shown to have potent SMARCA2 degrading activity in mouse tissue and xenograft tumors with robust tumor growth inhibitory activity SMARCA*4*-mutant xenograft tumors. Finally, we demonstrated synergistic combination effect of YDR1 with sotorasib against a sotorasib-resistant *KRAS^G12C^* and *SMARCA4* co-mutant cell line for the first time. We anticipate this study will stimulate future preclinical and clinical investigations to target *SMARCA4*-mutant cancers and expand the translational potential of SMARCA2 degraders as single or combination agents in the appropriate genomically defined subtypes of lung cancer.

## Author contributions

S.K. designed the studies, interpreted the data, wrote some sections of manuscript and performed most of the experiments with assistance from X.L., Y.W., Y.H, and N.B. Y.W, Y.J and P.K.N performed the *in vivo* pharmacology experiments. S.K. wrote the manuscript and formatted all figures. Y.L. conceived the work, designed the studies, designed PROTACs, oversaw the project and wrote the manuscript. All authors commented on the paper.

## Acknowledgements

We thank the MD Anderson Cancer Center (MDACC) Core facilities, including the histopathology core facility and Shan Jiang for assistance with mouse colony maintenance. We also thank Dr Jack Roth and other members of the Department of Thoracic Surgery-Research section for valuable comments during the progress of this project. This study was in part supported by the generous philanthropic contributions to The University of Texas MD Anderson Lung Cancer Moon Shots Program (YL). This work is also supported by NIH grants R01CA272945 (YL) and R37CA251629 (YL).

## Declaration of Interests

Y.L. is an inventor on a patent application, WO 2023/129506 A1, for YD54, YDR1 and related compounds. The authors declare no competing interests.

## Materials and Methods

### Chemical synthesis of YDR1 and YD54

All commercial chemical reagents and solvents were used directly without additional purification, unless specifically indicated. Liquid Chromatography Mass Spectra (LCMS) were recorded on SHIMADZU LCMS-2020 and High-Performance Liquid Chromatography (HPLC) were recorded on SHIMADZU LC-20AB. Proton nuclear magnetic resonance (^1^H NMR) spectra and (^19^F NMR) spectra were recorded on a Bruker 400 MHz instrument. Chemical shifts were reported in parts per million (ppm) and the residual solvent peak was used as an internal reference: proton (chloroform δ 7.27, DMSO δ 2.50, CH3OH δ 3.34). The NMR data was expressed in the following format: (d) chemical shift (multiplicity, J values in Hz, integration). The abbreviations used are as follows: s = singlet, d = doublet, t = triplet, q = quartet, m = multiplet. Most compounds were purified by column chromatography (SiO2). Purity of final compound was analyzed by HPLC analysis (SHIMADZU LC-20AB; XBridge C18, 2.1 X 50 mm,5 um; 10%–80% MeCN in H2O containing 0.025% NH3·H2O in 6 min; 0.8 mL/min). Synthesis and characterization of YDR1, YD54, and related compounds are detailed below. NMR Spectra for compounds YDR1 and YD54, and HPLC spectra for YDR1 and YD54 are provided (Supplemental Data 1 and 2).

Synthetic Scheme for YDR1

### Preparation of YDR1

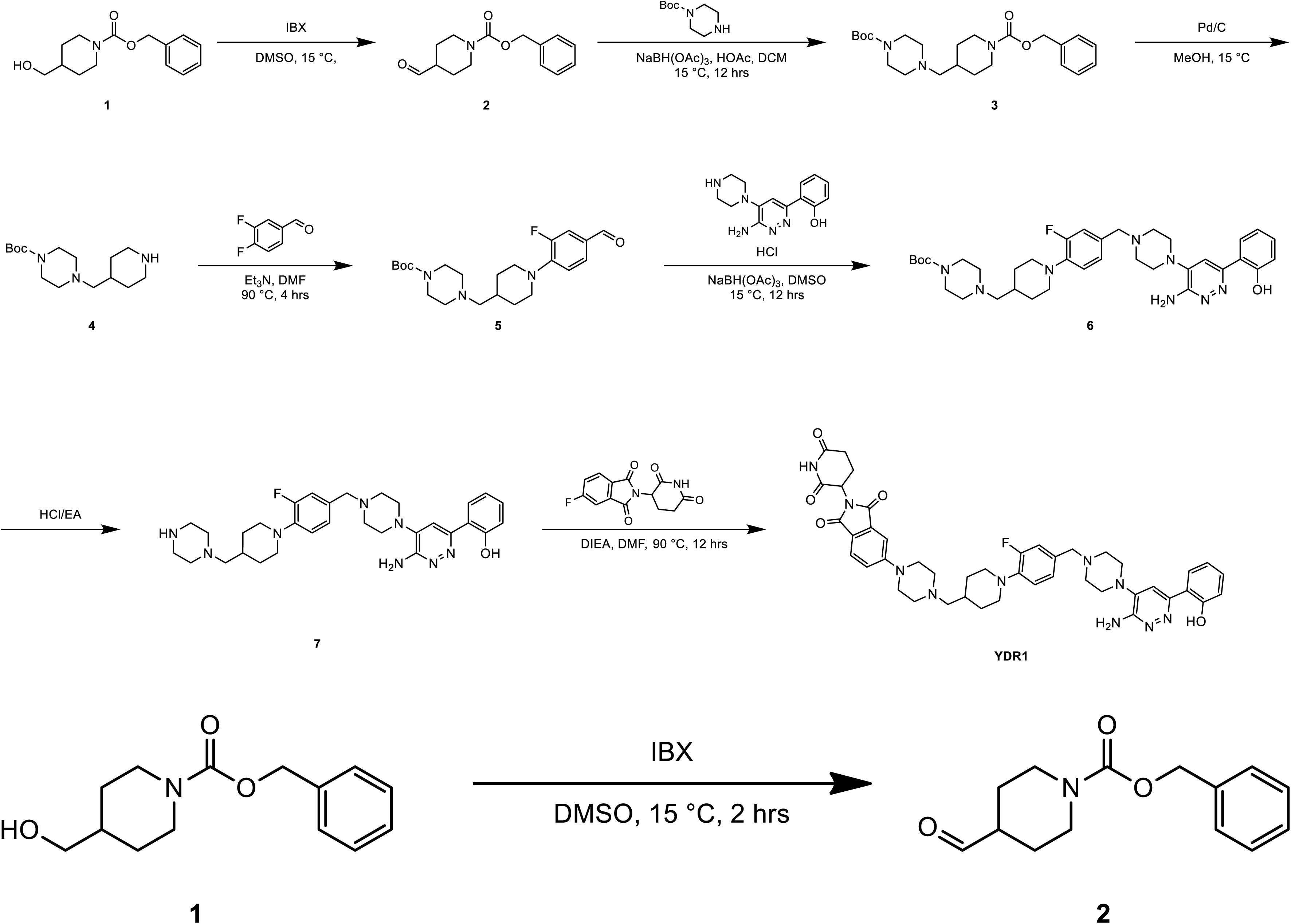

To a solution of **compound 1** (4.00 g, 16.0 mmol, 1.00 *eq*) in DMSO (40.0 mL) was added IBX (6.74 g, 24.1 mmol, 1.50 *eq*). The mixture was stirred at 15 °C for 2 hrs. LCMS (EW28004-8-P1A2) showed **compound 1** was consumed completely and one main peak with desired mass was detected. The mixture was diluted with H_2_O (100 mL) and extracted with EtOAc (100 mL × 2). The combined organic layers were washed with brine (100 mL), dried over Na_2_SO_4_, filtered and concentrated under reduced pressure to give a residue to afford the crude compound **2** (3.70 g, crude) as a colorless oil and was used into the next step without further purification.

LCMS: EW28004-8-P1B, R_t_ = 0. 993 min, *m/z* = 464.2 (M+1)

^1^H NMR: EW28004-8-P1A (400 MHz, CDCl_3_)

δ 9.67 (s, H), 7.39∼7.31 (m, 5H), 5.14 (s, 2H), 4.08∼4.05 (m, 2H), 3.07∼3.01 (m, 2H), 2.48∼2.41 (m, 1H), 1.91 (m, 2H), 1.64∼1.54 (m, 2H)

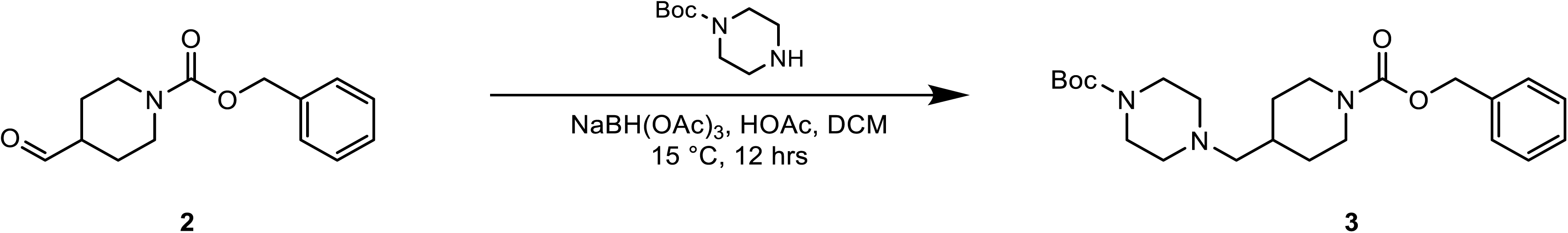

To a solution of **compound 2** (3.70 g, 14.9 mmol, 1.00 *eq*) and tert-butyl piperazine-1-carboxylate (2.93 g, 15.7 mmol, 1.05 *eq*) in (40.0 mL) was added HOAc (898 mg, 14.9 mmol, 856 uL, 1.00 *eq*) and NaBH(OAc)_3_ (4.76 g, 22.4 mmol, 1.50 *eq*). The mixture was stirred at 15 °C for 12 hrs. LCMS (EW28004-9-P1A) showed compound **2** was consumed completely and one main peak with desired mass was detected. The mixture was diluted with H_2_O (100 mL) and extracted with DCM (100 mL × 2). The combined organic layers were washed with brine (100 mL), dried over Na_2_SO_4_, filtered and concentrated under reduced pressure to give a residue to afford **compound 3** (6.00 g, crude) as a white solid.

LCMS: EW28004-9-P1B, R_t_ = 0.798 min, *m/z* = 418.2 (M+1)

^1^H NMR: EW28004-9-P1A (400 MHz, CDCl_3_)

δ 7.39∼7.30 (m, 5H), 5.13 (s, 2H), 4.18 (s, 2H), 3.41 (d, J = 4.8 Hz, 4H), 2.77 (m, 2H), 2.34 (d, J = 4.8 Hz, 4H), 2.17 (t, J = 7.2 Hz, 2H), 1.77∼1.74 (m, 2H), 1.46 (s, 9H), 1.27 (m, 1H), 1.17∼1.07 (m, 2H)

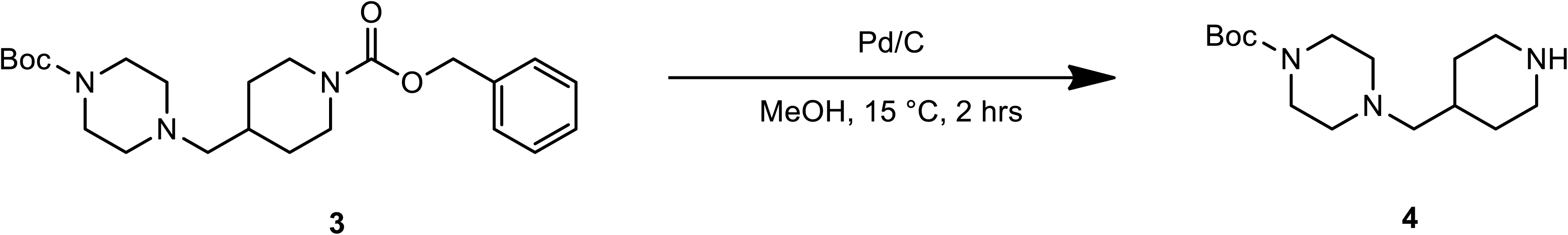

To a solution of compound **3** (4.00 g, 9.58 mmol, 1.00 *eq*) in MeOH (40.0 mL) was added Pd/C (0.40 g, 958 umol, 10.0 % purity, 0.10 *eq*) under N_2_. The suspension was degassed under vacuum and purged with H_2_ several times. The mixture was stirred under H_2_ (15 psi) at 15°C for 2 hrs. TLC (petroleum ether: ethyl acetate = 1: 1) showed Reactant 1 was consumed completely and one major new spot with larger polarity was detected. The mixture was filtered and the filtered cake washed with MeOH (100 ml), concentrated under reduced pressure to afford compound **4** (2.20 g, crude) as a gray solid.

^1^H NMR: EW28004-10-P1A (400 MHz, CDCl_3_)

δ 3.41 (d, J = 4.8 Hz, 4H), 3.10∼3.07 (m, 2H), 2.63∼2.51(m, 2H), 2.33 (d, J = 4.8 Hz, 4H), 2.16 (t, J = 7.2 Hz, 2H), 2.05 (s, 1H), 1.75∼1.72 (m, 2H), 1.65∼1.58 (m, 1H), 1.46 (s, 9H), 1.15∼1.05 (m, 2H)

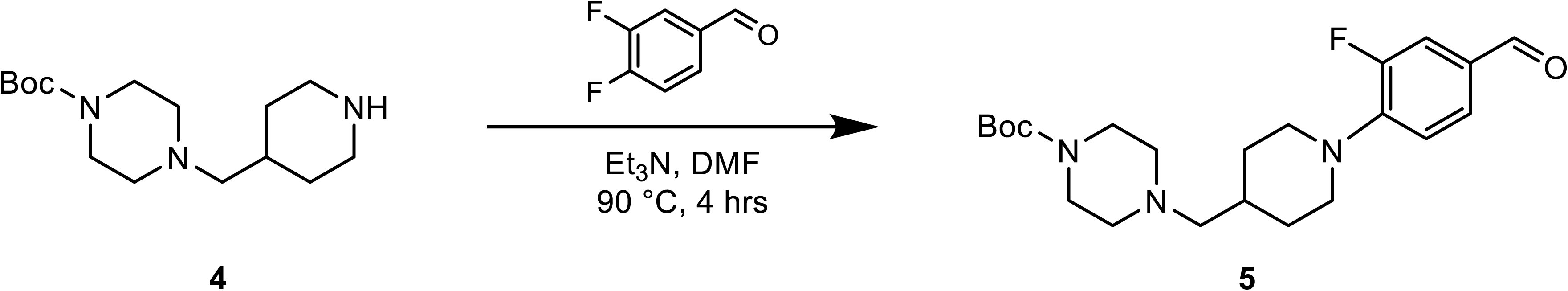

To a solution of compound **4** (2.20 g, 7.76 mmol, 1.00 *eq*) and 3,4-difluorobenzaldehyde (1.16 g, 8.15 mmol, 884 uL, 1.05 *eq*) in DMF (20.0 mL) was added Et_3_N (1.18 g, 11.6 mmol, 1.62 mL, 1.50 *eq*). The mixture was stirred at 90 °C for 4 hrs. LCMS (EW28004-11-P1A) showed compound **4A** was consumed completely and one main peak with desired mass was detected. The mixture was diluted with H_2_O (100 mL) and extracted with EtOAc (50 mL × 2). The combined organic layers were washed with brine (30 mL× 2), dried over Na_2_SO_4_, filtered and concentrated under reduced pressure to give a residue. The residue was purified by column chromatography (SiO_2_, Petroleum ether/Ethyl acetate=10/1 to 1/1, R_f_ = 0.40) to afford compound **5** (2.30 g, 4.82 mmol, 62.0% yield, 84.9% purity) as a yellow solid.

LCMS: EW28004-11-P1B, R_t_ = 0.799 min, *m/z* =406.2 (M+1) HPLC: EW28004-11-P1B1, R_t_ = 1.535 min

^1^H NMR: EW28004-11-P1A (400 MHz, CDCl_3_)

δ 9.81∼9.80 (m, 1H), 7.56∼7.48 (m, 2H), 6.99∼6.96 (m, 1H), 3.70∼3.67 (m, 2H), 3.43 (m, 4H), 2.85∼2.79 (m, 2H), 2.36 (m, 4H), 2.23 (t, J = 7.2 Hz, 2H), 1.88 (t, J = 12.4 Hz, 2H), 1.74∼1.68 (m, 1H), 1.47 (s, 9H), 1.40∼1.36 (m, 2H)

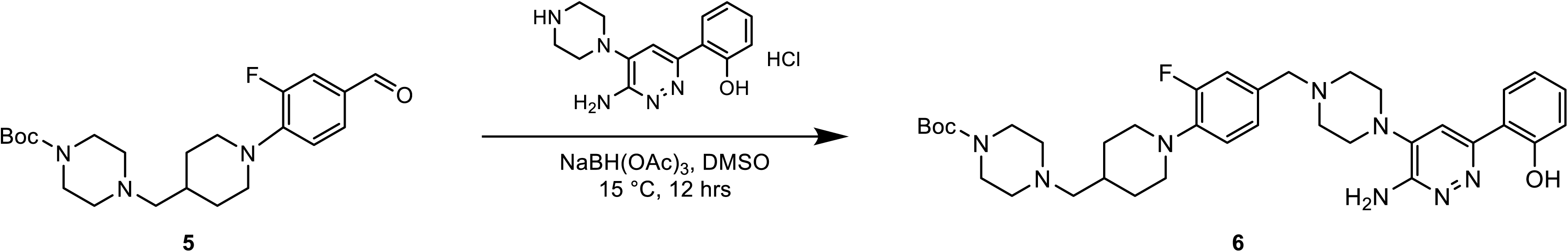

To a solution of **compound 5** (1.00 g, 2.47 mmol, 1.00 *eq*) and 2-(6-amino-5-(piperazin-1-yl)pyridazin-3-yl)phenol hydrochloride (704 mg, 2.59 mmol, 1.05 *eq*) in DMSO (10.0 mL) was added NaBH(OAc)_3_ (1.05 g, 4.94 mmol, 2.00 *eq*). The mixture was stirred at 15 °C for 12 hrs. LCMS (EW28004-12-P1A) showed **compound 5** was consumed completely and one main peak with desired mass was detected. The mixture was diluted with H_2_O (30 mL) and extracted with EtOAc (50 mL × 2). The combined organic layers were washed with brine (30 mL× 2), dried over Na_2_SO_4_, filtered and concentrated under reduced pressure to give a residue. The crude product was triturated with PE (10.0 mL) for 15 mins. Then the mixture was filtered and the filtered cake was washed with Petroleum ether (3 mL × 2) to afford compound **6** (1.00 g, 1.37 mmol, 55.3% yield, 90.3% purity) as a yellow solid.

LCMS: EW28004-12-P1B, R_t_ = 0.741 min, *m/z* =661.3 (M+1)

^1^H NMR: EW28004-12-P1A (400 MHz, CDCl_3_)

δ 13.7 (s, 1H), 7.59∼7.57 (m, 1H), 7.32∼7.28 (m, 2H), 7.06∼7.00 (m, 3H), 6.94∼6.89 (m, 2H), 4.85 (s, 2H), 3.54 (s, 2H), 3.45 (m, 6H), 3.17 (s, 3H), 2.69∼2.64 (m, 6H), 2.40 (s, 4H), 2.28∼2.26 (m, 2H), 1.89∼1.87 (m, 2H), 1.66 (m, 1H), 1.47 (s, 9H), 1.40∼1.36 (m, 2H)

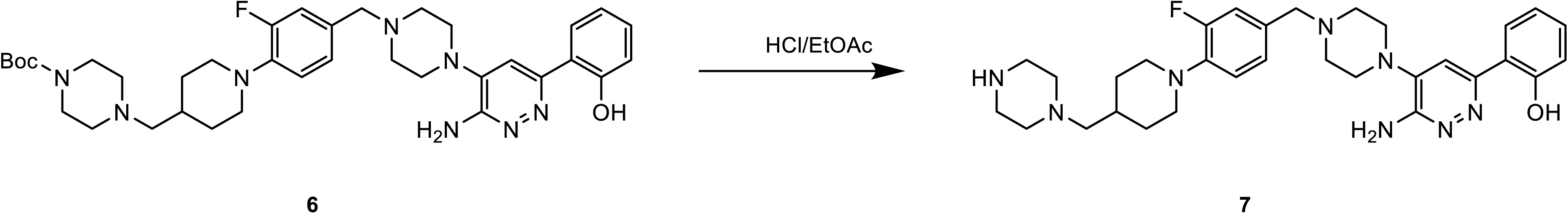

To a solution of **compound 6** (500 mg, 756 umol, 1.00 *eq*) in HCl/EtOAc (5.00 mL) was stirred at 15 °C for 2 hrs. LCMS (EW28004-13-P1A) showed **compound 6** was consumed completely and one main peak with desired mass was detected. The mixture was concentrated under reduced pressure to give a residue to afford compound **7** (400 mg, crude, HCl) as yellow solid.

LCMS: EW28004-13-P1B, R_t_ = 0.711 min, *m/z* =561.3 (M+1)

^1^H NMR: EW28004-13-P1A (400 MHz, CDCl_3_)

δ 12.7 (s, 1H), 10.2 (s, 2H), 7.59∼7.57 (m, 1H), 7.32∼7.28 (m, 2H), 7.06∼7.00 (m, 3H), 6.94∼6.89 (m, 2H), 4.85 (s, 2H), 3.54 (s, 2H), 3.45 (m, 6H), 3.17 (s, 3H), 2.69∼2.64 (m, 6H), 2.40 (s, 4H), 2.28∼2.26 (m, 2H), 1.89∼1.87 (m, 2H), 1.66 (m, 1H), 1.40∼1.36 (m, 2H)

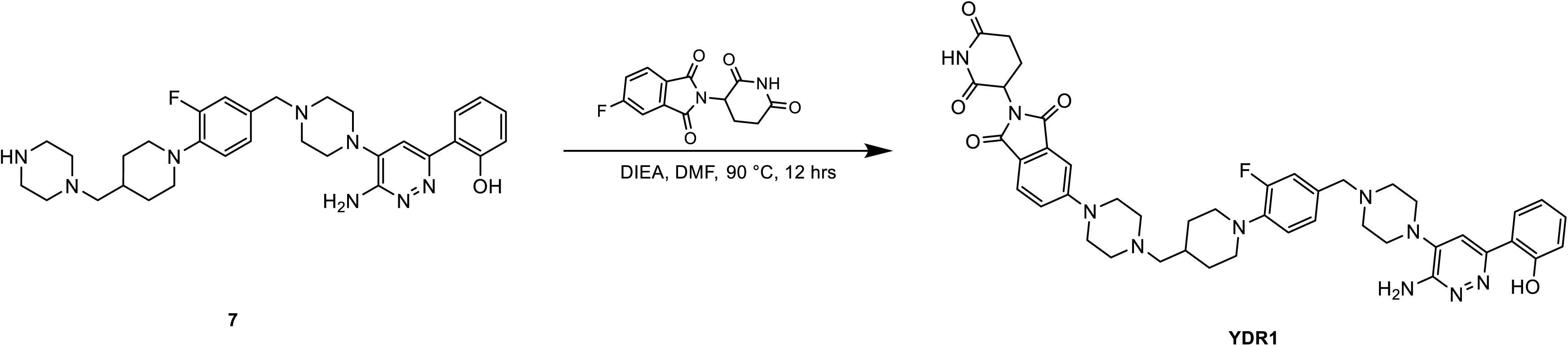

To a solution of **compound 7** (200 mg, 335 umol, 1.00 *eq*, HCl) and 2-(2,6-dioxopiperidin-3-yl)-5-fluoroisoindoline-1,3-dione (97.1 mg, 352 umol, 1.05 *eq*) in DMF (2.00 mL) was added DIEA (216 mg, 1.67 mmol, 292 uL, 5.00 *eq*). The mixture was stirred at 90 °C for 12 hrs. LCMS (EW28004-14-P1A3) showed **compound 7** was consumed completely and one main peak with desired mass was detected. The mixture was diluted with H_2_O (20.0 mL) and extracted with ethyl acetate (20.0 mL × 2). The combined organic layers were washed with brine (10.0 mL × 2), dried over Na_2_SO_4_, filtered and concentrated under reduced pressure to give a residue. The residue was purified by Prep-HPLC (column: Waters Xbridge C18 150*50mm* 10um;mobile phase: [water(10mM NH4HCO3)-ACN];B%: 48%-78%,11min) to afford **YDR1** (50.0 mg, 59.5 umol, 17.8% yield, 97.3% purity) as a yellow solid.

LCMS: EW28004-14-P1C1, R_t_ = 1.061 min, *m/z* =817.0 (M+1) HPLC: EW28004-14-P1C2, R_t_ = 2.611 min

^1^H NMR: EW28004-14-P1A (400 MHz, CDCl_3_)

**FNMR: EW28004-14-P1A**

δ 13.7 (s, 1H), 8.20 (s, 1H), 7.70 (t, J = 8.4 Hz, 2H), 7.60∼7.58 (m, 1H), 7.33 (s, 2H), 7.31∼7.29 (m, 2H), 4.97∼4.93 (m, 1H), 4.82 (s, 2H), 3.55 (s, 2H), 3.46∼3.40 (m, 6H), 3.18 (s, 4H), 2.92∼2.77 (m, 3H), 2.74∼2.72 (m, 1H), 2.71∼2.66 (m, 5H), 2.61 (s, 4H), 2.33 (t, J = 6.8 Hz, 2H), 2.18∼2.09 (m, 1H), 1.89 (t, J = 12.4 Hz, 2H), 1.53∼1.40 (m, 3H)

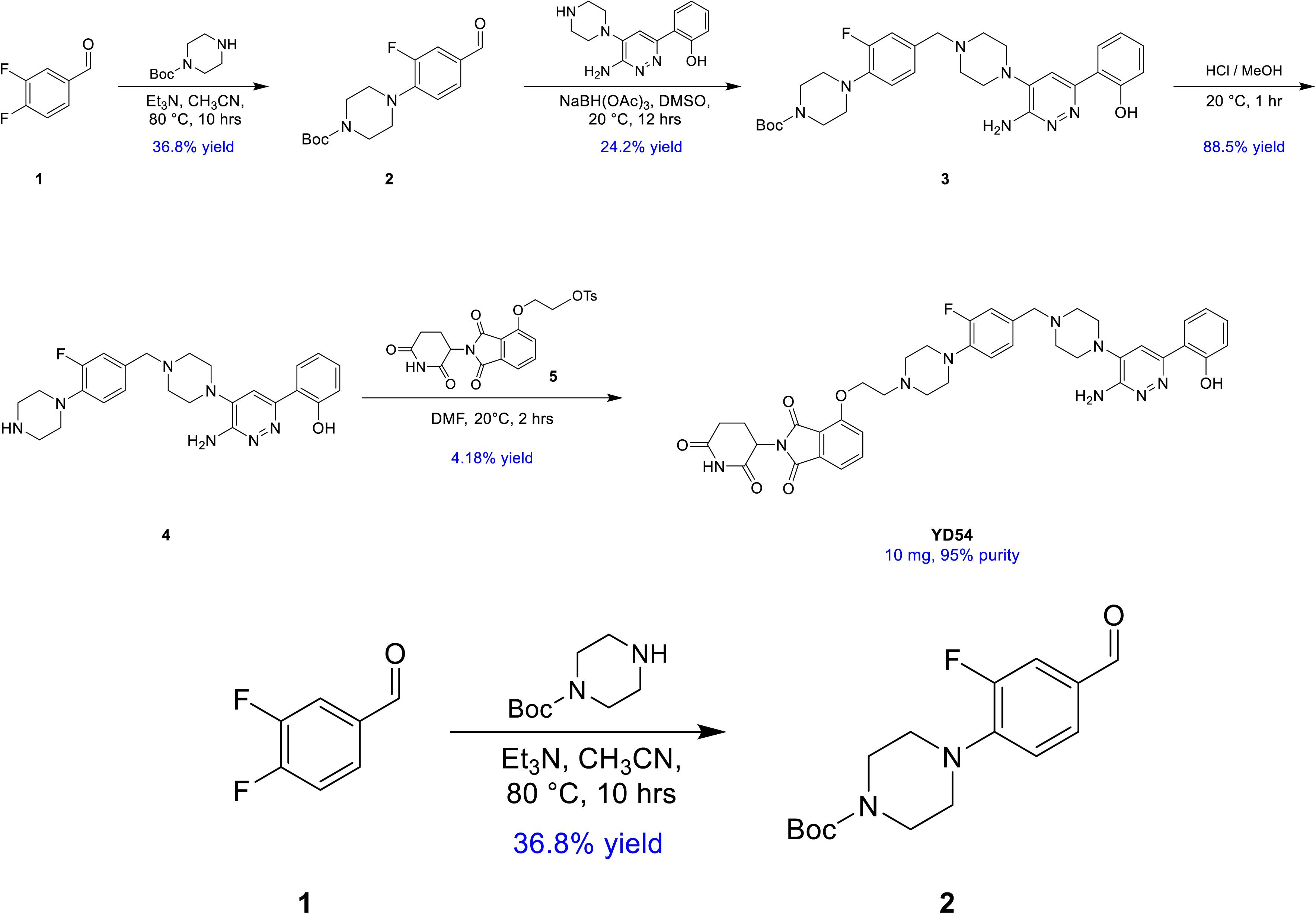

### Preparation of YD54

To a solution of **Cpd.1** (1.00 g, 7.04 mmol, 763 uL) in CH_3_CN (20.0 mL) was added Et_3_N (1.42 g, 14.0 mmol, 1.96 mL) and tert-butyl piperazine-1-carboxylate (1.57 g, 8.44 mmol). The mixture was stirred at 80°C for 10 hrs. TLC (Petroleum ether: Ethyl acetate=5:1) showed some of the starting material remained was and a main spot (Rf = 0.25) was formed. The reaction mixture was filtered and the filter liquor was concentrated to give the crude product. The residue was purified by column chromatography (SiO_2_, Petroleum ether/Ethyl acetate=30/1 to 5/1) to give **Cpd.2** (0.80 g, 2.59 mmol, 36.8% yield) as light yellow oil.

**LCMS:** EW16633-201-P1B1, R_t_ = 0.963 min, m/z = 253.1 (M+H)^+^

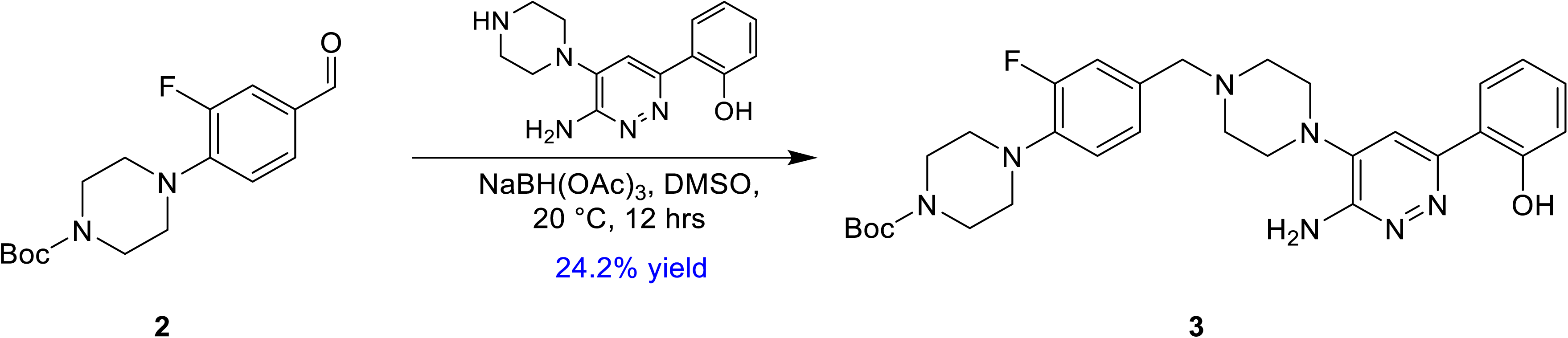

To a solution of **Cpd.2** (1.00 g, 3.24 mmol) in DMSO (10.0 mL) was added 2-(6-amino-5-piperazin-1-yl-pyridazin-3-yl) phenol (967 mg, 3.57 mmol, HCl salt), NaBH(OAc)_3_ (824 mg, 3.89 mmol) and Et_3_N (656 mg, 6.49 mmol, 902 uL). The mixture was stirred at 20°C for 12 hrs. TLC (Petroleum ether : Ethyl acetate=1:1,P1:Rf=0.25) showed the start material was remained and some new spots was detected. The reaction mixture was diluted with **H_2_O (30 mL)**, and then extracted with DCM (20 mL × **2**). The combined organic layers were washed with brine (10 mL*2), dried over Na_2_SO_4_, filtered and concentrated under reduced pressure to give a residue. The residue was purified by column chromatography (SiO2, Petroleum ether/Ethyl acetate=10/1 to 2/1) to afford **Cpd.3** (450 mg, 784 umol, 24.2% yield, 98.3% purity) as a white solid.

**LCMS:** EW20934-239-P1B3, R_t_ = 0.805 min, m/z = 564.4 (M+H)^+^

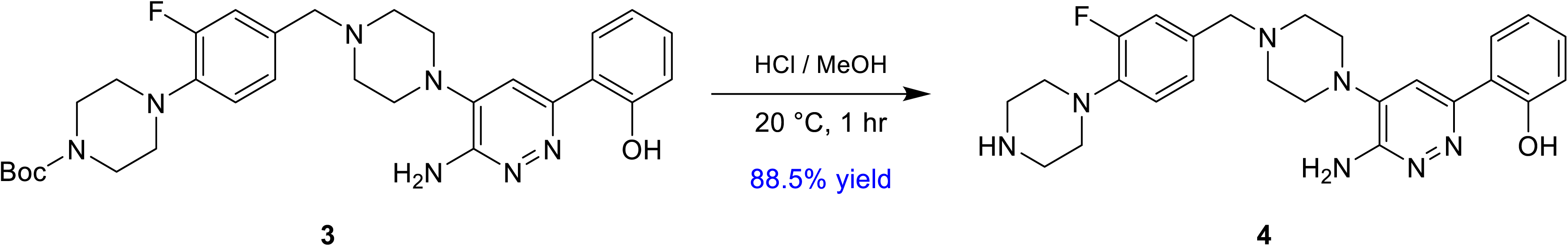

To a solution of **Cpd.3** (400 mg, 709 umol) in MeOH (4.00 mL) was added HCl/MeOH (4 M, 4.00 mL). Then the mixture was stirred at 20 °C for 1 hr. The mixture was concentrated under reduced pressure to give a residue. The mixture was suspended in DCM (10 mL) and then adjust pH to 8 – 9 with saturated NaHCO3 solution. Then the mixture was extracted with DCM (20.0 mL × 2). The combined organic layers were washed with brine (10 mL), dried over Na_2_SO_4_, filtered and concentrated under reduced pressure to afford **Cpd.4** (300 mg, 628 umol, 88.5% yield, 97.1% purity) as a white solid.

**LCMS:** EW20934-242-P1A3, R_t_ = 1.033 min, m/z = 464.1 (M+H)^+^

**^1^H NMR:** EW20934-242-P1A, (400 MHz, DMSO-*d6*)

*δ* 14.4 - 14.2 (m, 1H), 8.0 - 7.9 (m, 1H), 7.51 (s, 1H), 7.30 - 7.21 (m, 1H), 7.09 - 7.03 (m, 2H), 7.01 - 6.95 (m, 1H), 6.95 - 6.82 (m, 2H), 6.24 (s, 2H), 3.49 (s, 2H), 3.11 ( s, 4H), 3.02 - 2.98 (m, 8H), 2.58 (s, 5H)

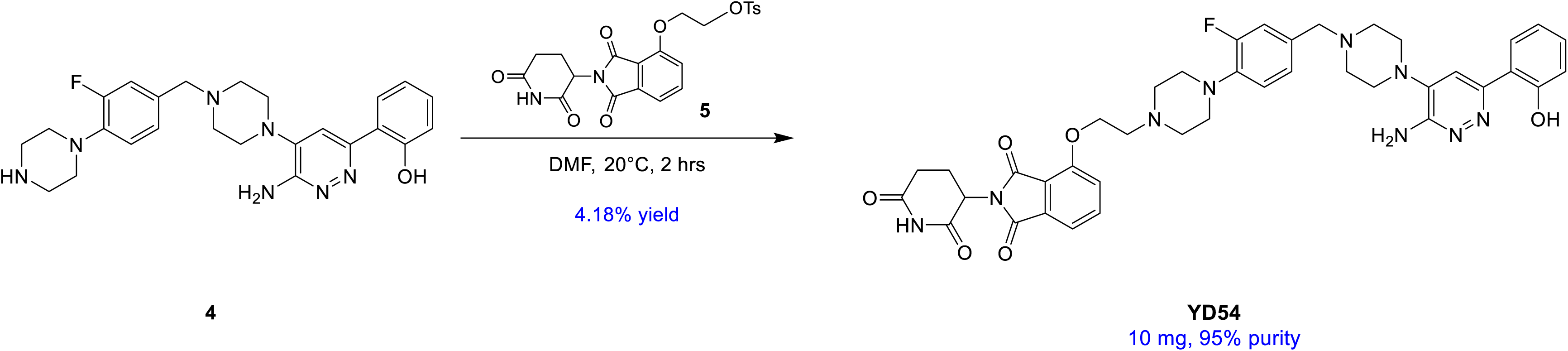

To a solution of **Cpd.4** (203 mg, 431 umol) in DMF (2.00 mL) was added **Cpd.5** (200 mg, 431 umol), Et_3_N (43.6 mg, 431 umol, 60.0 uL). The mixture was stirred at 20°C for 2 hrs. The mixtue was purified by Prep-HPLC (neutral condition) to give **YD54** (15.0 mg, 18.0 umol, 4.18% yield, 91.8% purity) as a white solid.

**LCMS:** EW28074-31-P1A1, R_t_ = 0.702 min, m/z = 764.3 (M+H)^+^

**HPLC:** EW28074-31-P1A2, R_t_ = 1.160 min, purity = 95.9%

^1^H **NMR:** EW28074-31-P1A (400 MHz, CDCl_3_)

*δ* 14.1 - 13.4 (m, 1H), 8.42 - 8.39 (m, 1H), 7.88 - 7.76 (m, 1H), 7.58 - 7.46 (m, 1H), 7.48 (m, 1H), 7.32 - 7.30 (m, 1H), 7.28 - 7.27 (m, 1H), 7.25 - 7.22 (m, 1H), 7.18 - 7.08 (m, 3H), 7.06 - 6.89 (m, 2H), 4.98 - 4.94 (m, 1H), 4.82 (s, 2H), 4.42 (s, 2H), 3.54 (s, 2H), 3.25 - 3.18 (m, 8H), 3.06 - 3.01 (m, 2H), 2.94 - 2.76 (m, 7H), 2.73 - 2.55 (m, 4H), 2.17 - 2.08 (m, 1H)

**F NMR:** EW28074-31-P1A (400 MHz, CDCl_3_)

**SFC:** EW28074-31-P1A_c1424, Rt= 0.555 min

#### Cell culture

H1792, H322, H2030, H2126, H1568, H2023 and H2030 cell lines were cultured in RPMI (Roswell Park Memorial Institute 1640 Medium, Hyclone, Cat# SH30027.01; no pyruvate) with 10% heat inactivated fetal bovine serum (Gibco, Cat# 1614007), 1% penicillin/streptomycin (Hyclone, Cat# SV30010), and 2 mM L-glutamine. Cell lines were cultured till 80% confluency and kept at low passage numbers and maintained at 37 °C in a humidified 5% CO2-containing incubator. H1792, H322, H2030, H2126, H1568, H2030, H2023 cells were obtained from ATCC. HCC-515 cell lines were obtained from University of Texas Southwestern Medical Centre. Routine Mycoplasma testing was performed using Mycoplasma Detection Kit (ABM, Cat # G238). STR profiling for cell lines were done at MD Anderson Cancer Center core facility.

#### Compounds and Antibodies

Methylcellulose (Cat# M0262) was purchased from Millipore sigma. Sotorasib (Cat# HY-114277) was purchased from MedChemExpress (South Brunswick Township, NJ). Antibodies against SMARCA2 (Cat# 11966, 1:1000) was from Cell Signaling (Danvers, MA, USA). β-actin (#A2228, 1:20,000) was from Millipore Sigma (Burlington, MA). **Cell proliferation and colony formation assays.** Single-cell suspensions of all cell lines were counted and plated into 6-well plates or 12-well plate at a density of 0.5–2 × 10^3^ cells per well. Cells were cultured in a medium containing the indicated drugs for 9–14 days (refreshed every 5^th^ day). At the endpoints, cells were fixed with Formalin (10%) and stained with crystal violet (0.01%w/v in water) and scanned. Total colony area was calculated with Gelcount (Optronix) instrument. All experiments were performed in triplicates. IncuCyte S3 (Essen Biosciences, Ann Arbor, MI, USA) was used to measure cell proliferation. For IncuCyte S3 experiments, H2030cells were suspended in fresh growth media, plated in 96-well plates at 500 cells/well, and grown overnight at 37 °C. On the following day, cells were treated with drug, and confluency was assessed every 4 hours till the end of the experiment. Confluency was determined at each time point using the IncuCyte S3 analysis software.

#### Immunohistochemistry

Xenograft lung tumors were fixed in formalin for 24 hours, paraffin embedded, sectioned, and stained according to standard procedures. After antigen retrieval in citrate buffer (pH: 6), slides were immersed in 3% hydrogen peroxide for 10 minutes. Non-specific signals were blocked for one hour using 2.5% normal goat serum. Slides were stained using respective antibodies overnight at 4°C, SMARCA2 (Cat#11966, Cel Signaling 1:2000). Slides were washed and incubated with ImmPRESS-HRP conjugated anti-mouse or anti-rabbit cocktail from Vector Laboratories (MP-7451 and MP-7452) for 30 min at room temperature. The slides were thrice washed and stained with DAB substrate (VECTOR, SK-4105). The slides were counterstained with hematoxylin for 1minute and mounted with mounting medium. After DAB staining, slides were dehydrated thrice in ethanol and thrice in xylene (1 min each), and mounted using Cytoseal 60. Images were further taken with the Vectra® 3 automated quantitative pathology imaging system. Five images were acquired for each tumor and images were processed using the Inform® 2.5 software from Akoya Biosciences®. Images were prepared based on a spectral library which detects DAB and Hematoxylin.

#### Immunoblotting

Cells were washed in ice-cold PBS, and proteins extracted in RIPA buffer having protease and phosphatase inhibitors for 30 minutes. Lysates were then collected and centrifuged at 14,000 rpm for 15 min at 4°C. Protein concentrations were measured using the BCA Protein Assay Kit (Thermo scientific). Lysates were mixed with freshly made 4X laemmli buffer, boiled at 95°C for 10 minutes and processed with Mini-Protean Gel Electrophoresis Systems (Bio-Rad) using Bis-Tris 4–20% gradient pre-cast gels (Biorad).

#### Calculation of DC_50_ and D_max_

The protein expression of SMARCA2, SMARCA4 and β-Actin was determined from western blots and ImageJ software was used for quantification. SMARCA2 and SMARCA4 expression was normalized to β-Actin. The expression level of SMARCA2 and SMARCA4 in DMSO-treated cells was defined as 100%. The expression of treated cells was defined as the percentage relative to DMSO-treated cells. Values were analyzed with GraphPad Prism software using non-linear regression analysis to determine DC_50_ and D_max_.

#### Pharmacokinetics

Male CD-1(ICR) mice were dosed intraperitoneally with vehicle, YDR1 or YD54 at 1.25 mg/mL in 5% DMSO / 5%Solutol / 90%Water, final pH=4.39 by pH meter, clear solution. NU/J nude mice were dosed oral gavage with vehicle (0.5% methylcellulose), YDR1 or YD54 at 1.25 mg/mL in 0.5% MC (4000 cps) / 0.2% Tween 80 in water, homogenous opaque suspension. Blood was collected to measure mean plasma concentration of YDR1 and YD54. All blank matrix was freshly prepared. Protein precipitation (PPT) was done using 96-well plate. An aliquot of 5 µL unknown sample, calibration standard, quality control and dilution quality control, single blank, and double blank sample was added to the 96-well plate respectively. Each sample (except the double blank) was quenched with 200 µL of IS1 respectively (double blank sample was quenched with 200 µL of ACN), and then the mixture was vortex-mixed for 10 min at 800 rpm and centrifuged for 15 min at 3220 × g, 4 °C. An aliquot of 50 µL supernatant was transferred to another clean 96-well plate and centrifuged for 5 min at 3220 × g, 4 °C, then the supernatant was directly injected for LC-MS/MS analysis.

### Liver microsomal assays

Working solutions of compound were prepared from 10mM stock solution in dimethyl sulfoxide (DMSO) by diluting with 495 μL of acetonitrile (ACN) for intermediate solution concentration of 100 μM, 99% ACN. Using an Apricot automation workstation, 2 μL/well of compound working solution were added to all 96-well reaction plates except the blank (T0, T5, T15, T30, T45, T60, and NCF60). Then, an Apricot automation workstation was used to add 100 μL/well of microsome solution to all reaction plates (Blank, T0, T5, T15, T30, T45, T60, and NCF60). All reaction plates containing mixtures of compound and microsomes were pre-incubated at 37°C for 10 minutes. An Apricot automation workstation was used to add 98 μL/well of 100 mM potassium phosphate buffer to reaction plate NCF60. Reaction plate NCF60 was incubated at 37°C for one hour. After pre-incubation, 98 μL/well of NADPH regenerating system was added to every reaction plate except NCF60 (Blank, T0, T5, T15, T30, T45 and T60) to start the reaction. Final concentration of each component in incubation medium (0.5mg protein/ml Microsome, 1μM test compound, 1μM control compound, 10μM ritonavir, 0.99%Acetonitrile, 0.01% DMSO). The reaction plates were incubated at 37°C. Six time points T0, T5,T15,T30,T45 and T60 minutes were included. An Apricot automation workstation was used to add 600 μL/well of stop solution (Cold (4°C) acetonitrile (ACN) containing 200 ng/mL tolbutamide and 200 ng/mL labetalol as internal standards) followed by NADPH solution to each reaction plate at its appropriate end time point to terminate the reaction. Each plate was sealed and shaken for 10 minutes. After shaking, each plate was centrifuged at 4000 rpm and 4°C for 20 minutes. After centrifugation, an Apricot automation workstation was used to transfer 300 μL of supernatant from each reaction plate to eight new 96-well plates to LC-MS/MS analysis. Cmax, T_1/2,_ and AUC are calculated for liver microsomal assays. Assays was performed with mouse liver microsome kit (Cat No. M1000), Xenotech.

### Human lung xenografts anti-tumor efficacy studies

Animal experiments were performed according to standards defined by following internationally recognized guidelines on animal welfare. Animal procedures were accepted by the Institutional Animal Care Committee according to guidelines of the MDACC. To establish xenograft models, H1568 cells (2×10^6^ cells/mouse), H2023 cells (10×10^6^ cells/mouse), were trypsinized and resuspended in 1X PBS, mixed with a 1:1 mix of Matrigel in a final volume of 200 μl, and injected subcutaneously into the flanks of nude female mice (Jackson Labs) at 6-8 weeks of age. Tumor volume was measured by external caliper. Mice were randomized to control and treatment groups once the average tumor volume of H1568 xenograft reached 60mm3 and H2023 xenograft tumor reached < 200mm^3^, depending on the lung model. Vehicle, YDR1 and YD54 was prepared in 0.5% Methylcellulose + 1% Tween-80 solution by sonication. Briefly, freshly prepared solutions were subjected to six rounds of sonication: each pulse of 15 seconds with a gap of 2 minutes, while on ice. Experimental mice were treated with vehicle, YDR1 or YD54 by oral gavage once daily until the end of the experiment. Tumor volume and mouse body weight were measured alternate days. Tumor diameter and volume were calculated based on caliper measurements of tumor length and height using the formula tumor volume = (length × width^2^)/2. The maximal tumor size permitted by our Institutional Animal Care and Use Committee is 2500 mm^3^.

#### Quantitative mass spectrometry

*SMARCA4*-WT, H1792 cell line was treated with YDR1 48 hr and submitted for multiplexed quantitative mass spectrometry analysis by Thermo Fisher Scientific Center for Multiplexed Proteomics (Harvard Medical School) processed and analyzed as previously described^58^. Sample processing steps included cell lysis, protein precipitation, tandem digestion with LysC and trypsin, peptide labeling with Tandem Mass Tag 6-plex reagents and off-line bRP HPLC fractionation. Multiplexed quantitative mass spectrometry data was collected on an Orbitrap Fusion mass spectrometer operating in a MS3 mode using synchronous precursor selection for the MS2 to MS3 fragmentation^59^. MS/MS data were searched against a Uniport human database (February 2014) with SEQUEST using a target-decoy search strategy^60^. Data processing steps included controlling peptide and protein level false discovery rates, assembling proteins from peptides, and protein quantification from TMT reporter ions as previously described^58^.

### Data mining and analysis

CRISPR/Cas9 knockout screening data were downloaded from the DepMap Public 23Q2 dataset (https://depmap.org/portal/). Cell lines were called as *SMARCA4*-WT when SMARCA4 is free of mutation and *SMARCA4*-mutant based on damaging mutations of *SMARCA4*.

#### Statistics and reproducibility

GraphPad Prism 9 software was used to generate graphs and statistical analyses. Statistical significance was determined by unpaired Student’s t-test. Methods for statistical tests, the exact value of n, and definition of error bars were showed in figure legends. All experiments have been reproduced in at least two independent experiments unless otherwise specified in the figure legends. All immunoblots and images shown are representatives of these independent experiments.

## Notes

### Competing Interest Statement

The authors have declared no competing interest.

